# Oncogenic memory underlying minimal residual disease in breast cancer

**DOI:** 10.1101/2020.01.23.916510

**Authors:** Ksenija Radic Shechter, Eleni Kafkia, Katharina Zirngibl, Sylwia Gawrzak, Ashna Alladin, Daniel Machado, Christian Lüchtenborg, Daniel C. Sévin, Britta Brügger, Kiran R. Patil, Martin Jechlinger

## Abstract

Tumor relapse is responsible for most breast cancer related deaths^1,2^. The disease recurrence stems from treatment refractory cancer cells that persist as minimal residual disease (MRD) for years following initial therapy^3^. Yet, the molecular characteristics defining the malignancy of MRD remain elusive due to difficulties in observing these rare cells in patients or in model organisms. Here, we use a tractable organoid system and multi-omics analysis to show that the dormant MRD cells retain metabolic peculiarities reminiscent of the tumor state. While the MRD cells were distinct from both normal and tumor cells at a global transcriptomic level, their metabolomic and lipidomic profile markedly resembled that of the tumor state. The MRD cells particularly exhibited a de-regulated urea cycle and elevated glycolysis. We find the latter being crucial for their survival and could be selectively targeted using a small molecule inhibitor of glycolytic activity. We validated these metabolic peculiarities of the MRD cells in corresponding tissues obtained from the mouse model as well as in transcriptomic data from patients following neo-adjuvant therapy. Together, our results show that the treatment surviving MRD cells retain features of the tumor state over an extended period suggestive of an oncogenic memory. In accord, we found striking similarity in DNA methylation profiles between the tumor and the MRD cells. The distinction of MRD from normal breast cells comes as a surprise, considering their phenotypic similarity with regards to proliferation and polarized epithelial organization. The metabolic aberrances of the MRD cells offer a therapeutic opportunity towards tackling emergence of breast tumor recurrence in post-treatment care.

---

Minimal residual disease (MRD) is estimated to lead to mostly incurable relapse in 20**-**40% of breast cancer patients within a period from a few years up to decades after the initial treatment^2,3^. Therefore, understanding and tackling MRD has been recognized as one of the great challenges to improve treatment options for breast cancer survivors^4,5^. MRD is not accessible for direct functional analysis in breast cancer patients due to its covert nature, but can be explored in mechanistic detail in model systems, such as genetically modified mouse models^6^. To this end, we have established a preclinical mouse model of breast cancer (see Methods) that bears doxycycline-inducible hMyc- and Neu/Her2-oncogenes^7–9^, which we have successfully used to isolate MRD relevant for human disease^10^. From these transgenic mice we obtain primary mammary cells to establish 3D organoid cultures^11^ and are able to follow MRD establishment in phenotypic and molecular detail. In short, polarized structures lining a lumen represent the healthy tissue (Fig. 1a, left panel, Normal), while addition of doxycycline at a dose of 200ng/ml (Extended data 1a-d) activates the expression of oncogenes, as shown for transgenic oncogenic-hMYC presence (Fig. 1a, right panels, Tumor). This leads to uncontrolled proliferation as well as the loss of polarity and lumen (Fig. 1a, left panels, Tumor). Removal of doxycycline from the media silences oncogene expression and triggers tumor regression, in line with the principle of oncogene dependence^12,13^. Residual acini exhibit a re-polarized epithelium and absence of hMYC protein (Fig. 1a, left and right panels, Residual), reminiscent to findings in mammary gland histological sections of the mouse model taken eight weeks after inducing tumor regression^10^.

**Figure 1:**
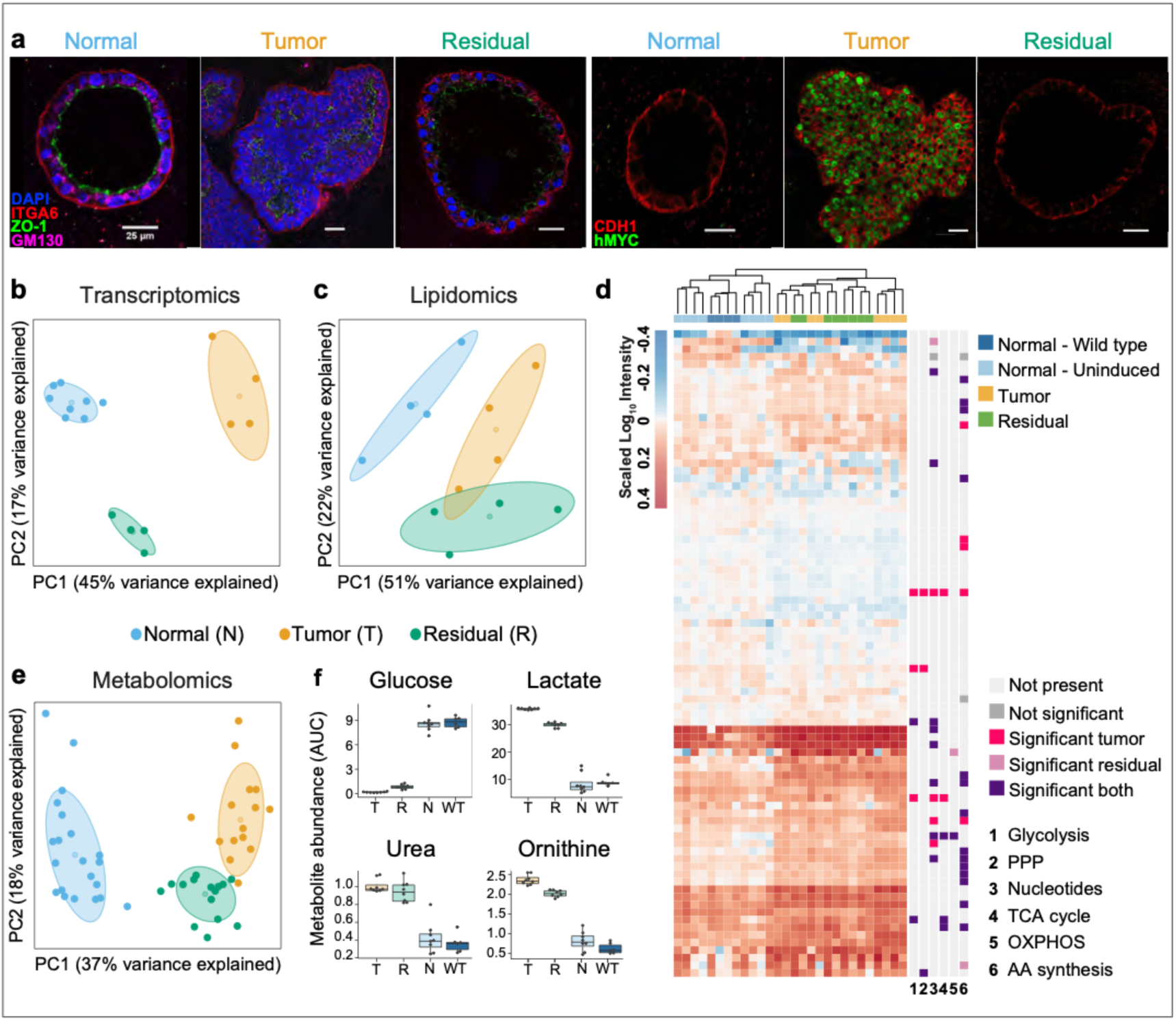
Multi-omics approach reveals characteristic features of residual structures *in vitro* and their metabolic resemblance to the tumor. **a**, Immunofluorescence staining of normal, tumor and residual structures. Left panel: polarity markers ITGA6 (red), ZO-1 (green), GM-130 (magenta); DAPI (blue). Right panel: human MYC oncogene (green); CDH1 (red). Scale bar: 25 μm. **b, c**, PCA analysis of normal (blue), tumor (orange) and residual (green) populations based on RNA sequencing data (b, normal n=8, tumor and residual n=4) and Lipidomics data (c, n=4). **d**, Heat map of untargeted metabolomics results showing the clustering of the three populations along with the most altered metabolic pathways (n=4). Hierarchical clustering was based on the complete linkage method using the Euclidean distance metric. Significance thresholds correspond to a *p*-value < 0.05 calculated using unpaired two-sided t-tests and adjusted for multiple hypothesis testing ^27^ in comparison to the normal population. PPP, pentose phosphate pathway; TCA, tricarboxylic acid cycle; OXPHOS, oxidative phosphorylation; AA amino acid. **e**, PCA analysis of normal and WT (blue), tumor (orange) and residual (green) populations based on intracellular metabolomics measurements targeted at central carbon metabolism (n=8; WT n=2). **f**, Selection of the most profoundly altered metabolites in extracellular spent growth media of normal (N and WT, n=4 and n=2, respectively), tumor (T) and residual (R) populations (n=4). The results are based on a metabolomics analysis targeted at central carbon metabolites. The values represent metabolite abundance levels as quantified by the area under the curve (AUC) of the corresponding ions for each metabolite. **a**-**f**, Number of replicates corresponds to different animals. **b**, **c**, **e**, Centroids represent the mean and concentration ellipses one standard deviation (level=0.68) of an estimated t-distribution based on the first two principal components. **f**, Box plots: midline, median; box, 25–75th percentile; whisker, minimum to maximum. Statistics are calculated using the limma package^28^ in R with the significance threshold corresponding to a Benjamini-Hochberg adjusted *p*-value ≤ 0.01 and a log2 fold change (residual or tumor compared to normal) ≥ 1 or ≤ -1.

Despite the phenotypic similarity of the residual spheres to normal acini, RNA-sequencing data showed that the residual cells exhibit a distinct transcriptional profile (Fig. 1b, Extended data 2a). Gene-set enrichment analysis of differentially expressed genes in residual cells in comparison to normal cells resulted in Gene Ontology (GO) terms of “cell division and cell cycle”, “cell signaling”, and “response to stimuli” being enriched for downregulated genes and are in line with the observed dormancy of the residual cells (Extended data 2b, Supplementary table 1). Genes connected to “cytoskeletal localization”, “cell adhesion and movement” and “activation of cell surface receptor signaling” were upregulated, supporting the re-polarization process upon MRD establishment (Extended data 2c). Additionally, the growth supporting metabolic pathways of pentose phosphate pathway and glycolysis, were enriched for significantly upregulated genes and were found similarly upregulated in tumor cells (Extended data 2d, bottom-Supplementary table 2). Sub-setting the transcriptome to metabolic genes, the distinct PCA distribution patterns of the three populations observed in the global transcriptome were retained (Extended data 2e), suggesting altogether a potential metabolic abnormality of the surviving cells.

To examine the metabolic state of the three cell populations in detail, we embarked on lipidomic profiling as well as untargeted and targeted metabolomics analyses. Both intracellular and extracellular (culture supernatant) samples were analyzed to obtain a comprehensive overview of the metabolic (patho)physiology. We first optimized the metabolite extractions from 3D cultures (Extended data 3). Then we obtained profiles from shotgun lipidomics, which showed a surprisingly close similarity of residual and tumor populations (Fig. 1c). This was also evident in the untargeted metabolomics analysis, with the regressed cells resembling tumor state and not the normal (Fig. 1d). Importantly, control samples obtained from wild-type mice lacking the *reverse tetracycline-controlled transactivator* (rtTA), but treated with doxycycline, clustered with the normal controls, thus excluding a potential confounding effect of doxycycline on the metabolism. Metabolomics analysis targeted at central carbon metabolism confirmed the results obtained from lipidomics and untargeted metabolomics, at the same time attesting that the dormant residual cells still retained key characteristics of their past tumorigenic state (Fig. 1e, Extended data 4a, Extended data 5). Notably, the residual cells aligned with the tumor in one of the universal cancer metabolic features, an enhanced glycolytic phenotype, as evident in decreased glucose pool sizes concomitant with increased levels of lactic acid (Fig. 1f). Another evident metabolic alteration prominent in the residual as well as in tumorigenic cells included an enhanced urea cycle activity shown by increased secretion of urea and ornithine (Fig. 1f).

As changes in metabolite pools do not necessarily reflect flux changes, we used an integrative genome-scale metabolic modelling approach combining transcriptomics and extracellular metabolomics data with flux-balance analysis. In addition, we also performed reporter metabolite analysis^14^ which integrates differential gene expression into the metabolic network to identify de-regulated metabolites (suggestive of changes in their turnover rate) (see Methods). Both genome-scale flux estimates and the reporter metabolite analysis corroborate the metabolite measurements confirming the upregulation of glycolysis and the urea cycle as major hallmarks of the residual cells. De-regulation of the urea cycle was confirmed by measured changes in other metabolites related to this pathway, including putrescine, proline, fumarate and aspartate (Extended data 5), with the latter two bridging also to the TCA cycle and nucleotide metabolism (Fig. 2, Extended data 6). Furthermore, the data also brought forward a de-regulation of the pentose phosphate pathway, the TCA cycle, glutamine uptake and S-Adenosyl methionine (SAM) metabolism (Fig. 2, Extended data 6). Together, the transcriptomics and metabolomics data, as well as flux modelling reveal significant metabolic peculiarities in the residual cell population reminiscent of the tumor state (compare Extended data 6 and 7).

**Figure 2:**
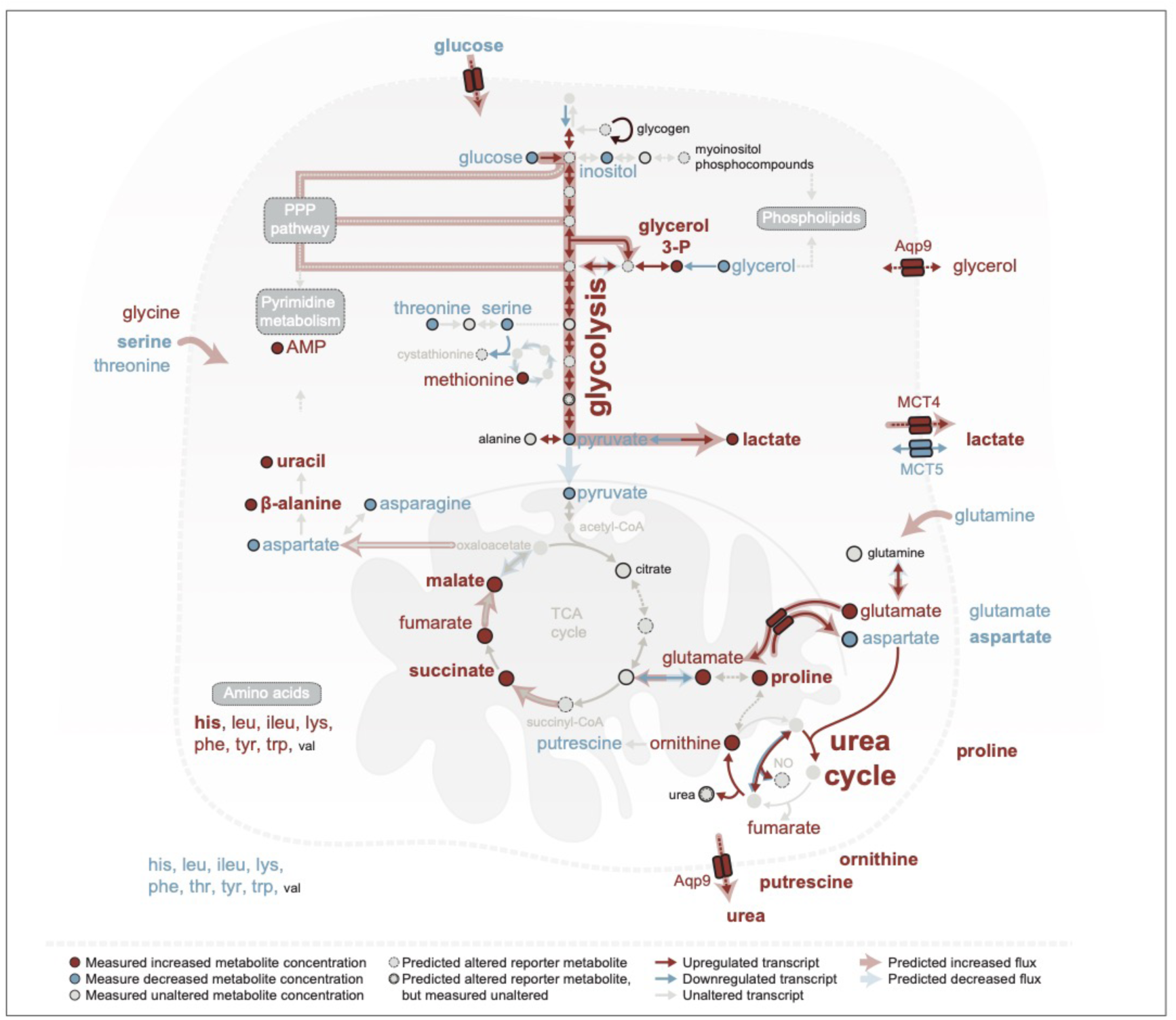
Global overview of altered metabolic pathways in residual cells, compared to normal. A selection of genes with significantly altered expression (Wald test^29^, Bonferroni adjusted *p*-value < 0.1; normal n=8, tumor and residual n=4), targeted metabolites with significantly altered levels (Benjamini-Hochberg adjusted *p*-value < 0.01; n=8; WT n=2) and significant reporter metabolites (top 5% with *p*-value < 0.1) of core metabolic processes are presented. Metabolites with an additional log_2_ fold change ≥ 1 or ≤ -1 are highlighted in bold. A Wald test with a Negative Binomial GLM was used as test statistics for gene expression and for metabolite levels. For the reporter metabolites a gene set enrichment analysis was performed from a theoretical null-distribution using the reporter method^30^. Bonferroni adjusted *p*-values of the expression analysis were used as gene-level statistics. Metabolite-gene sets were derived from a genome wide human metabolic model (HMR2), with genes mapped to mouse orthologs^31^. Flux balance analysis predicted metabolic fluxes, which are altered in the corresponding pathways, are overlaid. Significantly altered gene expressions and metabolites were used to inform the predictions. Number of replicates corresponds to different animals.

We next asked to what extent these metabolic features of residual cells in the organoid system compare with the situation in mice and patients. Mouse mammary glands taken nine weeks following oncogene in-activation and subsequent successful tumor regression were compared to healthy glands obtained from age-matched control animals. Indeed, metabolic changes still persisted in the regressed tissue (Fig. 3a) as reflected in elevated extracellular levels of urea, ornithine and putrescine (Fig. 3b), a higher percentage of ARG1 positive cells (Fig. 3c, Extended data 8), higher levels on NO (Fig. 3d) and higher flux to lactate (Fig. 3e). To assess the occurrence of such metabolic alterations in human patients, transcriptomic datasets of patient breast tissues after neoadjuvant treatment, were compared to breast tissues of healthy women. Notably, treated patient samples --based on genes in KEGG pathways being deregulated in cancer and involving HER2--clustered closely (Fig. 3f) with the residual mouse samples (presented in Fig. 1b) and showed similar alterations in glycolysis and urea cycle compared to controls (Fig. 3g). Additionally, alterations in these two pathways were also seen in the group of patient samples that were classified HER2 positive by histological analysis in the clinic (Extended data 9a) as well as in samples that clustered closely to mouse when the joint clustering was done based on all genes generally involved in HER2 or MYC related KEGG pathways (Extended data 9b, c). Taken together, these results support that changes in metabolism could be of potential relevance in clinical treatment^15^, which is specifically interesting for glycolysis^16,17^ and gives a working rational to test and re-purpose already approved metabolic drugs designed for primary tumor intervention^18^.

**Figure 3:**
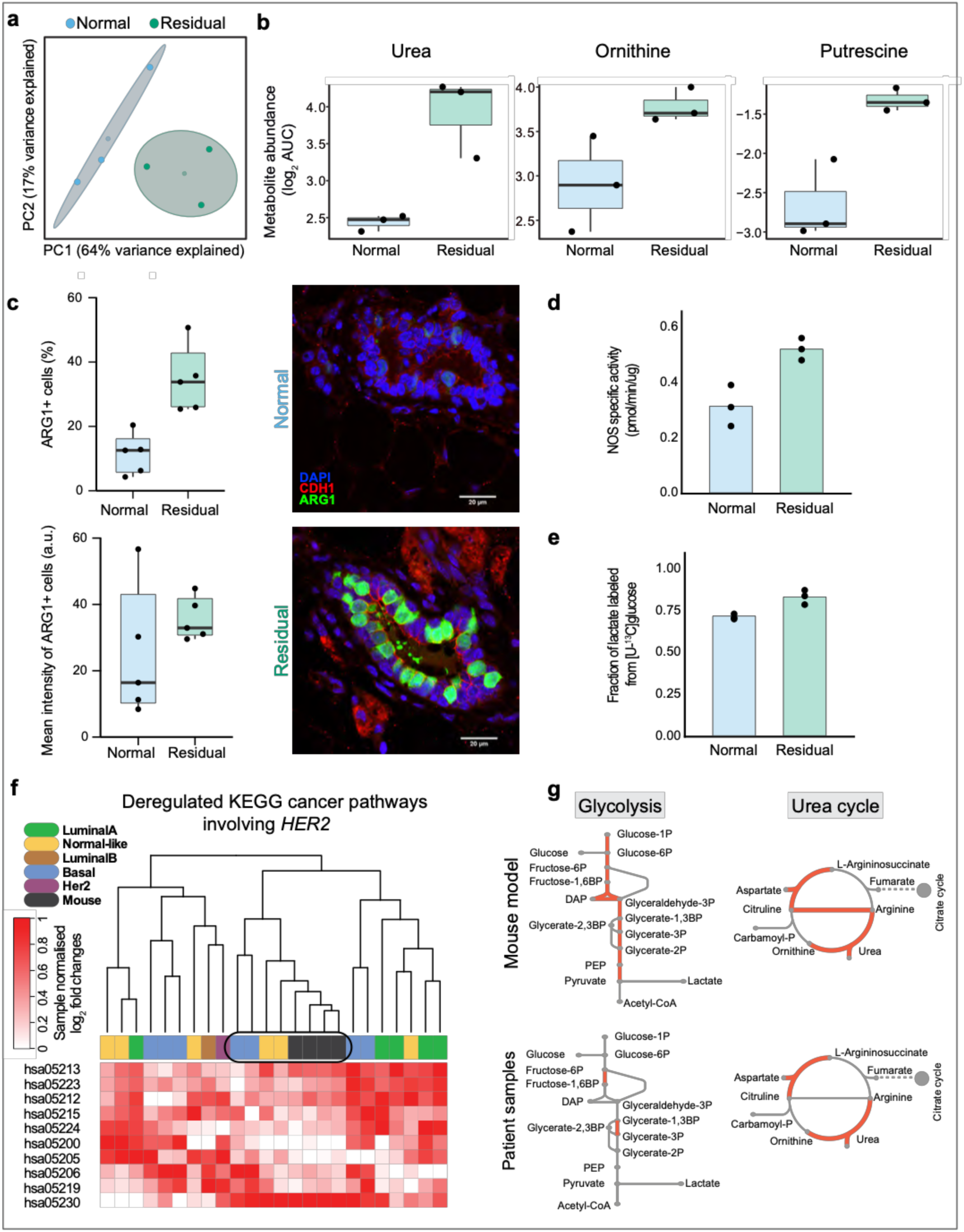
Glycolysis and urea cycle are the main altered metabolic pathways in residual cells in mice and human datasets. **a**, PCA analysis of extracellular metabolic profiles of isolated healthy (n=3; blue) and regressed (n=3; green) mammary glands following cultivation in cell growth media for 8 hours. The metabolomics analysis is targeted to central carbon metabolites. Centroids represent the mean and concentration ellipses one standard deviation (level = 0.68) of an estimated t-distribution based on the first two principal components. **b**, Selective secreted metabolites linked to urea cycle from healthy (n=3; blue) and regressed (n=3; green) mammary glands. The values represent metabolite abundance levels as quantified by the area under the curve (AUC) of the corresponding quantifying ions for each metabolite. **c**, Quantified ARG1+ cells (top) and intensity of ARG1 (bottom) in mammary gland sections of normal (n=5, 2921 cells) and residual (n=5, 2241 cells). Representative images of immunofluorescence staining in normal (top) and residual (bottom) duct stained for ARG1 (green); CDH1 (red), DAPI (blue). Scale bar: 20 μm. **d**, NOS activity in healthy (n=3) and regressed (n=3) mammary glands. **e**, Fractional labeling of lactate following cultivation of isolated regressed (n=3) and healthy (n=3) mammary glands in cell growth media supplemented with [U-^13^C] glucose for 8 hours. The three-carbon labeled (^13^C) isotopologue (M+3) is depicted. **f**, Joint clustering of mouse model (RNA-seq, normal n=8, residual n=4) and patient (microarray, healthy n=10 samples, residual n=20 samples) derived transcriptome log_2_ fold changes of healthy samples compared with regressed tumor samples. The clustering is based on all genes of KEGG pathways, which involve *HER2* and are known to be deregulated in cancer. Hierarchical clustering with the complete linkage method and the Euclidean distance as a distance metric was used for clustering. For the patient comparison, two independent data sets, one obtained from healthy breast tissue (GSE65194)^32,33^ and one obtained from patient tissues after neoadjuvant treatment (GSE32072)^34^, were merged. **g**, Metabolic reactions of glycolysis and urea cycle, which are catalyzed by enzymes with differential expression (Benjamini-Hochberg adjusted *p*-value < 0.1), are highlighted in red. An empirical Bayes moderated t-statistics was computed from a gene wise linear model fit with generalized least squares^28^, comparing the treated patients (n=4 samples), which are clustering closely with the mouse samples (**f**), with healthy breast tissue (n=10 samples). Differential expression from mouse *in vitro* transcriptome data of residual versus normal samples (RNA-seq, normal n=8, residual n=4) is shown in comparison (Wald test, Bonferroni adjusted *p*-value < 0.1)^29^. **a**-**f**, Number of replicates corresponds to different animals, **b**-**e**, Box plots: midline, median; box, 25–75th percentile; whisker, minimum to maximum. **b**, Statistics are calculated using the limma package^28^ in R with the significance threshold corresponding to a Benjamini-Hochberg adjusted *p*-value ≤ 0.01 and a log_2_ fold change (regressed or tumor compared to healthy) ≥ 1 or ≤ -1. **d**, **e**, The difference is statistically significant by unpaired two-samples t-test with *p*-values equal to 0.0136 and 0.0115, respectively.

Therefore, we examined whether inhibition of glycolysis was affecting the robustness of the residual population. Indeed, treatment with 3-bromopyruvate (3-BP), a well-established inhibitor of glycolysis^19^ induced cell death in residual cells grown in the absence of oncogenes for 10 days (Fig. 4a, Extended data 10a, b). This cytotoxic effect was especially prominent at the dose of 50μM (within the common range used for tissue culture^20,21^) and also reflected in the morphology of the residual cell structures, but not of normal acini (Fig. 4b, Extended data 10b). Increased cell death upon 3-BP treatment was also observed in the residual cells that were allowed to grow for 21 days in the absence of oncogenes expression (Fig. 4c), reinforcing the notion of a “memory” carried over from the tumor state. Quantification of the extracellular metabolite pools revealed that the relative fold change in glucose concentration upon treatment (3-BP to control glucose ratio) was the most dramatic for the regressed population, especially at the 50- and 250 µM doses (Fig. 4d), suggestive of a reduced glucose uptake. Consistently, beyond glucose, we additionally observed significant alterations in lactate pools and in glutamine levels in regressed cells following treatment (Extended data 10c). In comparison, normal cells were affected by 3-BP only to a minor extent. Together, the differences in cell death and glucose levels confirm the glycolytic nature of the residual cell population and its relevance as a targetable vulnerability.

**Figure 4:**
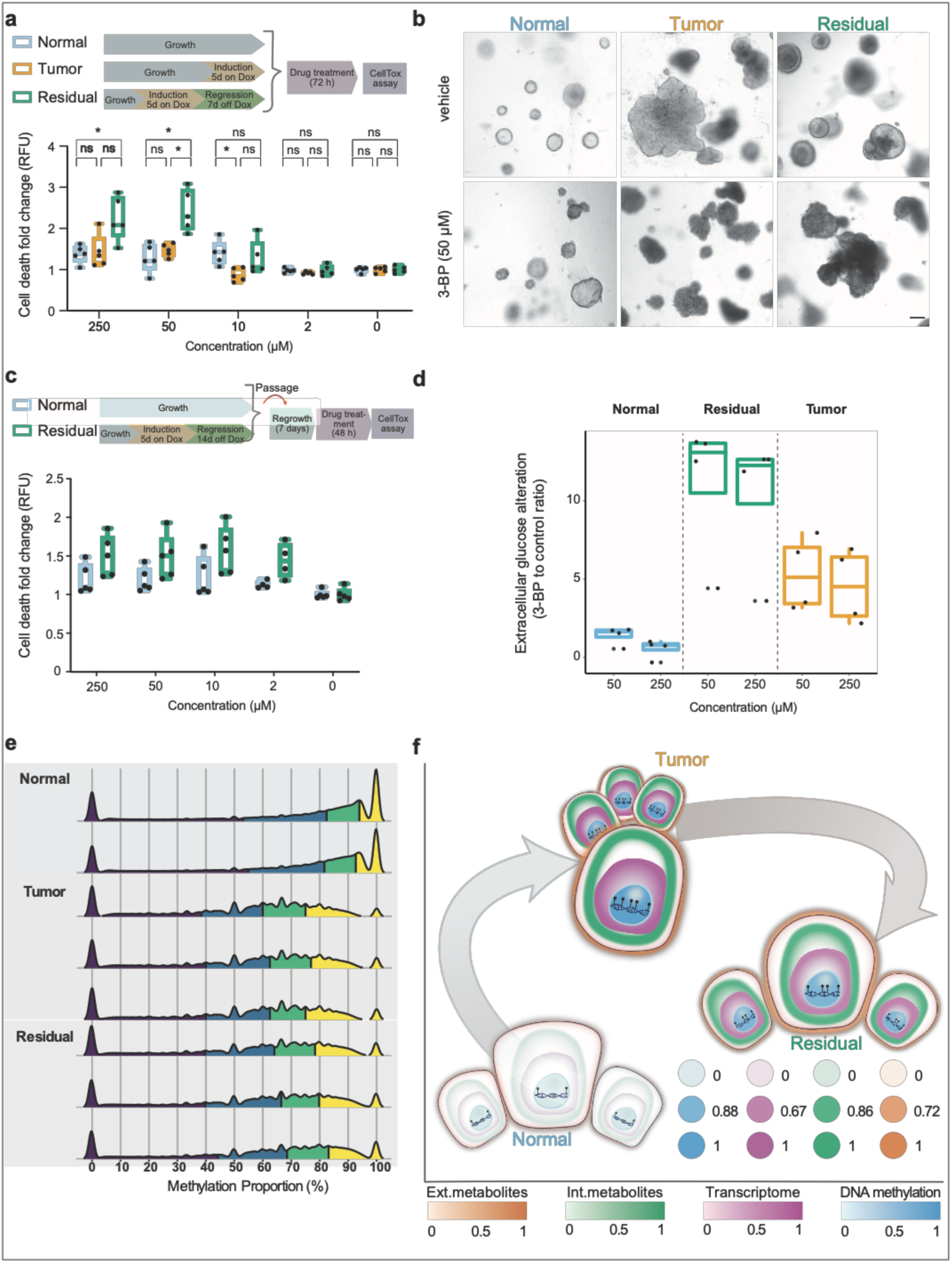
Residual cells require altered glycolysis for survival and maintain a DNA methylation profile similar to the tumor cell population. **a**, Experimental design (top) and cell death quantification in normal, tumor and residual structures after 72-hour treatment with 3-BP at the indicated doses (bottom) (n=5). **b**, Bright field images of healthy (left), tumor (middle) and residual (right) structures, treated with vehicle (top) and with 50 μM 3-BP (bottom), scale bar 100 μm. **c**, Experimental design (top) and cell death quantification in passaged residual and normal structures 72-hour treatment with 3-BP (n=5). **d**, Extracellular glucose abundance alteration upon treatment with 3-BP (at doses of 50- and 250 µM) in all three populations (n=4). The values represent the glucose ratio of 3-BP treated to untreated cells. **e**, DNA methylation profiles of normal (n=2), tumor (n=3) and residual (n=3) cell structures. Colors of the density plots represent quartiles. **f**, Summary figure integrating transcriptomics (normal n=8, tumor and residual n=4), intracellular metabolomics (n=8; WT n=2), extracellular metabolomics of spent growth media (normal n=4, WT n=2, tumor and residual n=3) and DNA methylomics (normal n=2, tumor and residual n=3) from the three populations. The color depth represents the normalized Euclidean distance of the respective omics layer in reference to normal. The distances between the centers of the three populations correspond to the normalized mean Euclidean distances across all represented omics layers. **a**-**f**, Number of replicates corresponds to different animals. **a**, **c**, **d**, Box plots: midline, median; box, 25–75th percentile; whisker, minimum to maximum.

As shown above, residual cells *in vitro* as well as MRD retain metabolic changes acquired during the tumor stage. Hinting at the mechanistic basis for this “oncogenic memory”, we find that the residual cells resemble tumor cells also in their DNA methylation profiles (Fig. 4e). Interestingly, certain metabolites that we found to be increased in the tumorigenic and residual populations (Extended data 5), like succinate, fumarate and nitric oxide, are also implicated in epigenetic modifications through direct inhibition of alpha-ketoglutarate dependent demethylases^22,23^. Additionally, succinate, lactate and nitric oxide, all accumulated in tumor and residual cells, can operate as signaling molecules by binding and stabilizing proteins that further interfere with hypoxic signaling, including the stabilization of hypoxia inducible factor 1 (HIF1α), a well-known master regulator of the glycolytic phenotype in cancers^24–26^. Overall, the persistence of a tumor-associated metabolic signature in the residual population, despite the absence of continued oncogenic input, suggests that the accumulation of certain metabolites offers an additional survival advantage for MRD that is sustained through epigenetically-imprinted signaling pathways.

Taken together, our study offers a first in depth characterization of MRD derived from a primary organoid system that is relevant for human disease^10^. These findings can be validated *in vivo* by directly analyzing the residual mammary glands taken from the mouse model (Fig. 3a-e). The residual cells, despite being characterized by a non-proliferative and histologically normal phenotype, harbor metabolic aberrations similar to that of the tumor cells. We refer to these changes as “oncogenic memory”, which is reflected in metabolite levels, glycolytic flux and DNA methylation patterns, and persists in the face of an inactive oncogenic signal in residual cells. Since residual cells represent a treatment refractory population of dormant cancer cells, tackling these memorized cancer hallmarks offers a unique therapeutic opportunity. We consequently illustrate how the interference with one of these metabolic nodes, viz., glycolysis, specifically targets the survival of residual cells. We envisage that the identified transcriptional, metabolic and epigenetic distinctiveness (Fig. 4f) of MRD will offer novel long-term treatment options for targeted interference to block the progression towards tumor relapse.

**Extended data 1:**
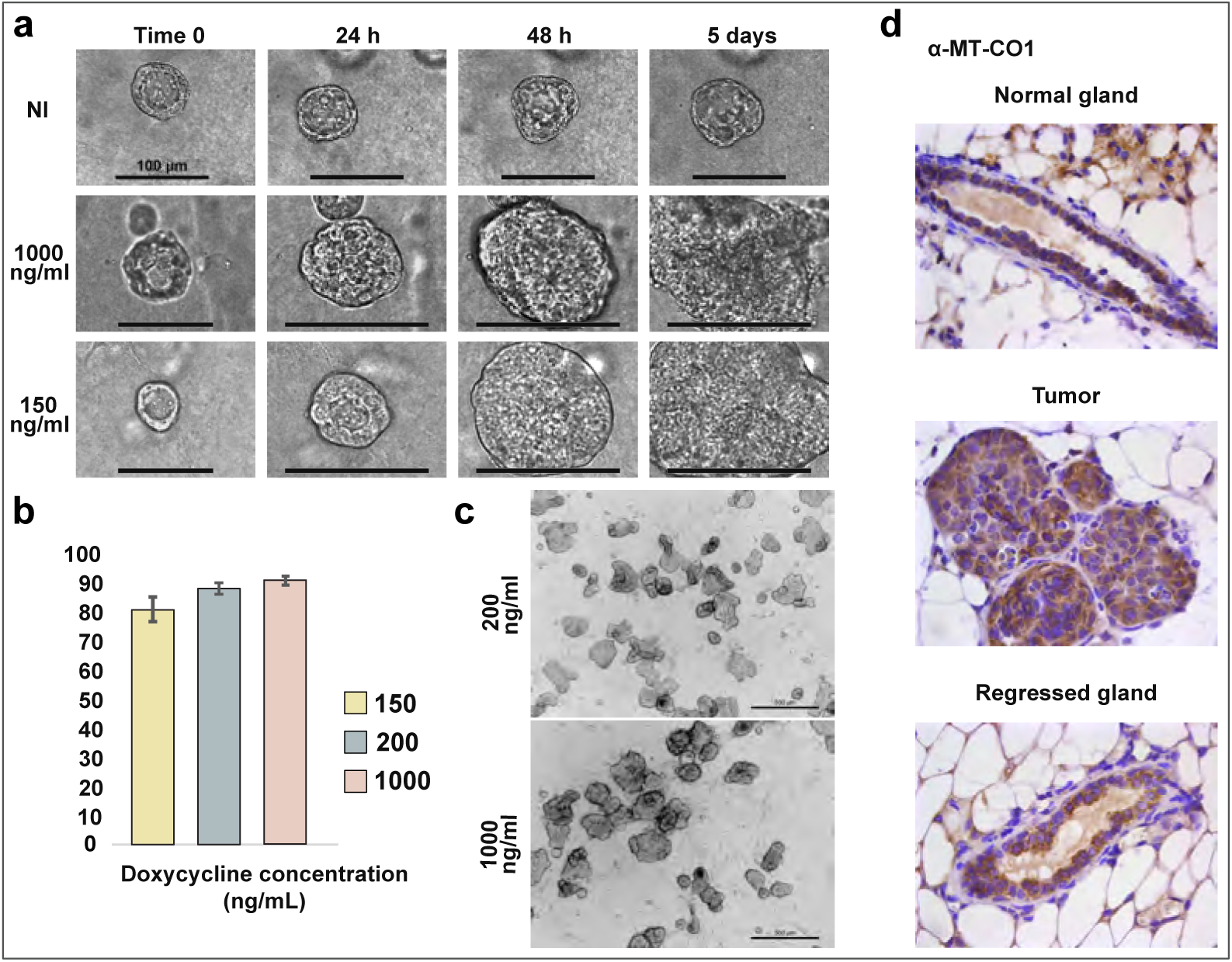
Effect of doxycycline concentration on phenotype, tumor induction and mitochondrial function. **a**, Bright field images of 3D cultures at indicated time points and doxycycline concentrations. Scale bar: 100 μm. **b**, Percentage of induced structures *in vitro* (5th day on doxycycline) at the concentrations of 150 (n=2), 200 (n=2) and 1000 (n=4) ng/mL. Data shown as mean ± SEM. **c**, Bright field images of induced structures on the 5th day on doxycycline at the concentration of 200 ng/mL (top) and 1000 ng/mL (below). Scale bar: 500 μm. **d**, IHC on MT-CO1 protein *in vivo*. From left to right: gland from age-matched control, tumor at 3 weeks on doxycycline, regressed gland at 2 weeks off doxycycline. Magnification: 4x. **a**-**d**, Number of replicates corresponds to different animals.

**Extended data 2:**
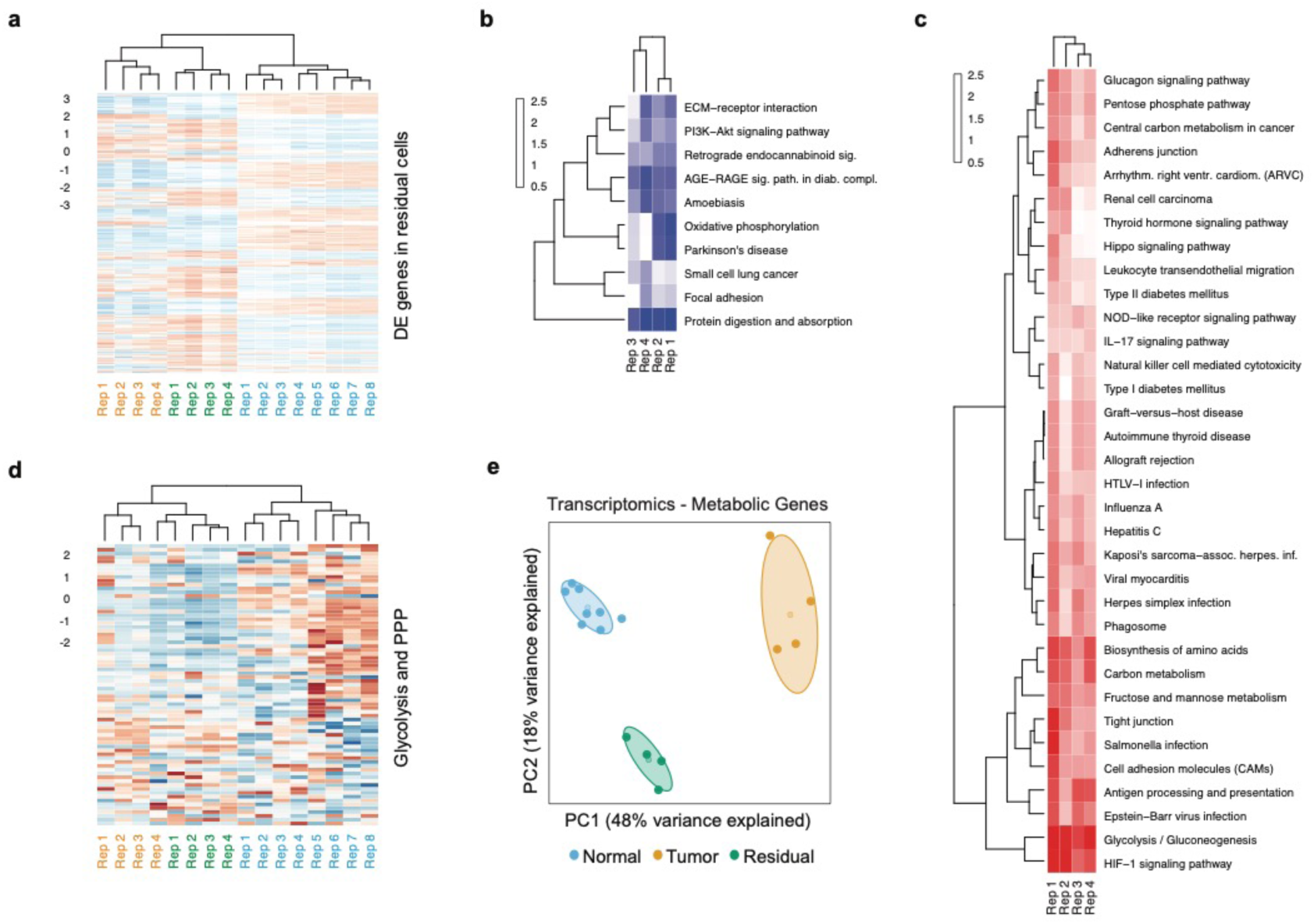
Transcriptome data analysis of different gene subsets. **a**, Heatmap with rlog transformed and gene-wise centered and scaled (stdv = 1) transcript counts of differentially expressed genes (Wald test^29^, Bonferroni adjusted *p*-value < 0.1) in residual cells compared to normal. **b**-**c**, Heatmap of significantly enriched KEGG pathways (unpaired two-sample t-test, *p*-value < 0.05) from differentially **b**, down-regulated and **c**, upregulated genes (Wald test, Bonferroni adjusted *p*-value < 0.1) in residual cells compared to normal. Clustering is based on fold changes of the sample wise calculated test statistics for the individual KEGG pathways of residual cells compared to normal. **d**, Heatmap with rlog transformed and gene-wise centered and scaled (stdv = 1) transcript counts of genes from glycolysis and pentose phosphate pathway. **e**, PCA analysis of transcriptome data sub-setted to metabolic genes, extracted from the mouse orthologs associated with the genome wide human metabolic model HMR2^31^. Centroids represent the mean and concentration ellipses one standard deviation (level=0.68) of an estimated t-distribution based on the first two principal components. **a**-**e**, Number of replicates (normal n=8, tumor and residual n=4) corresponds to different animals. **a**-**d**, Hierarchical clustering with the complete linkage method and the Euclidean distance as a distance metric was used for clustering. **a**, **d**, **e**, Sample labels: normal – blue, tumor – yellow, residual – green.

**Extended data 3:**
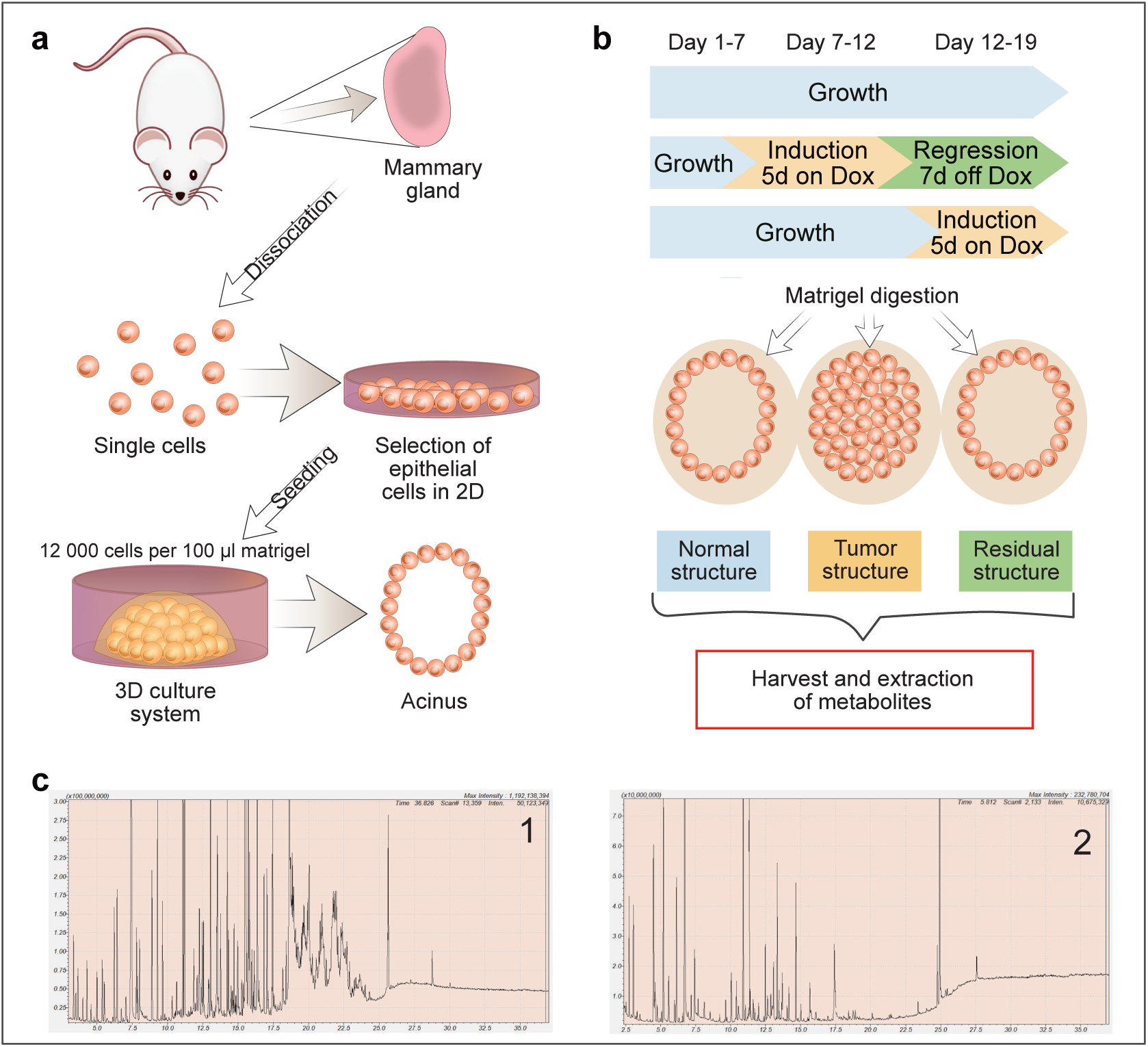
Set-up of metabolic methods for 3D cultures. **a**, Primary epithelial cells obtained from mouse mammary glands are grown in matrigel, which enables growth in 3D and formation of hollow acini, schematic **b**, Timeline of experiment showing period of growth, tumor induction (5 days on doxycycline) and tumor regression (after doxycycline withdrawal, 7 days off). Experiments were designed in such a way that the metabolites from all three conditions (normal, tumor and residual) were harvested at the same endpoint after matrigel digestion. **c**, Total ion chromatograms showing a strong matrigel signal when no digestion is performed for the collection of structures (1), and an improved signal of intracellular metabolites after gel digestion (2), representative example.

**Extended data 4:**
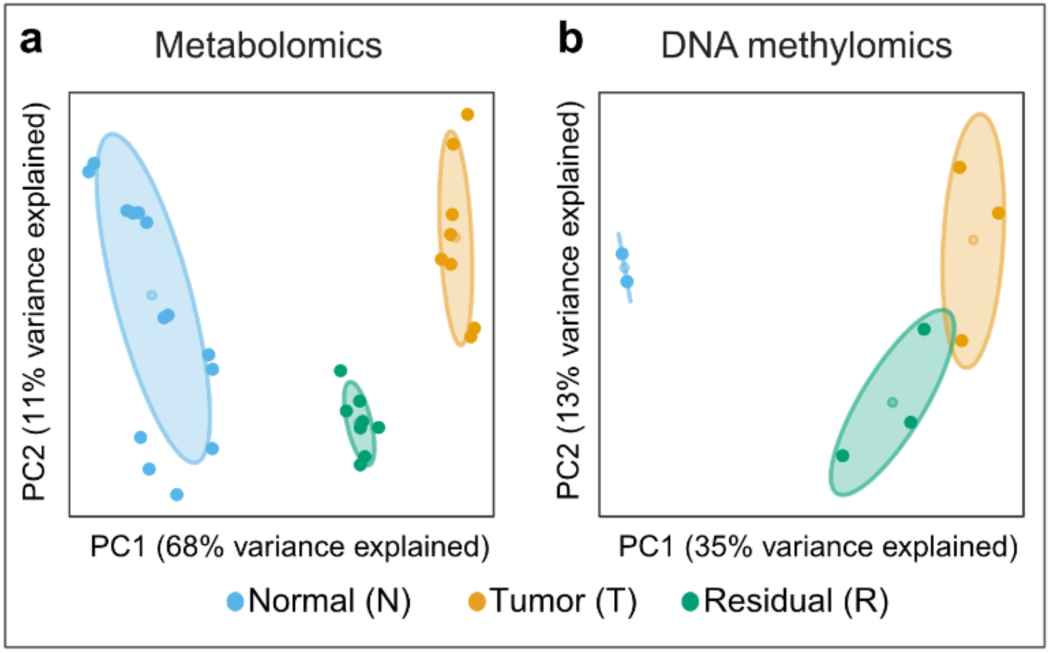
PCA analysis of additional omics layers revealing the residual structures’ resemblance to the tumor. **a**, PCA analysis of normal (blue; n=4, WT n=2), tumor (orange; n=3) and residual (green, n=3) populations based on the extracellular spent growth media targeted at central carbon metabolites. Centroids represent the mean and concentration ellipses one standard deviation (level=0.68) of an estimated t-distribution based on the first two principal components. **b**, PCA analysis of normal (blue; n=2), tumor (orange; n=3) and residual (green; n=3) populations based on genome wide DNA methylation data. The ellipses are confidence ellipses with 0.99 normal probability and the centroids represent the geometric center of the ellipse. This ellipse type was chosen because of the low number of replicates. **a**, **b**, Number of replicates corresponds to different animals.

**Extended data 5:**
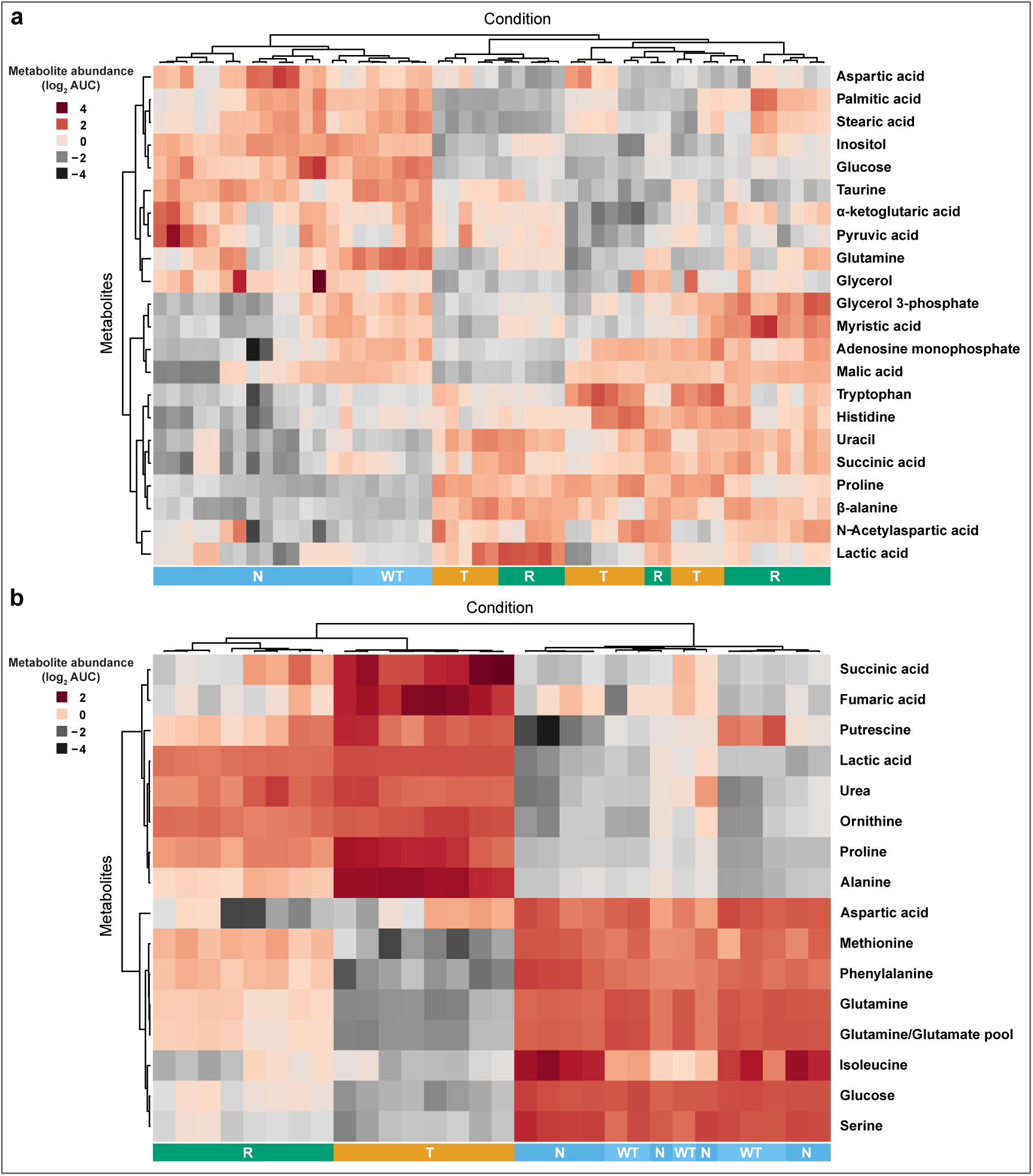
Residual population exhibits metabolic resemblance to the tumor. Heat maps representing the significantly altered metabolites in intracellular samples **(a)** and extracellular spent growth media **(b)** of residual (R, n=4) and tumor (T, n=4) compared to normal (N, n=4; WT n=2) cells. The results are based on metabolomics analyses targeted at central carbon metabolites. The hierarchical clustering of the samples and the metabolites is based on Pearson’s correlation. Statistics are calculated using the limma package^28^ in R with the significance threshold corresponding to a Benjamini-Hochberg adjusted *p*-value ≤ 0.01 and a log_2_ fold change (tumor or residual compared to normal cells) ≥ 1 or ≤ -1. Number of replicates corresponds to different animals. expression analysis (Wald test) were used as gene-level statistics. Metabolite-gene sets were derived from a genome wide human metabolic model (HMR2), with genes mapped to mouse orthologs^31^. Flux balance analysis predicted metabolic fluxes, which are altered in the corresponding pathways, are overlaid. Significantly altered gene expressions and metabolites were used to inform the predictions. Number of replicates corresponds to different animals.

**Extended data 6:**
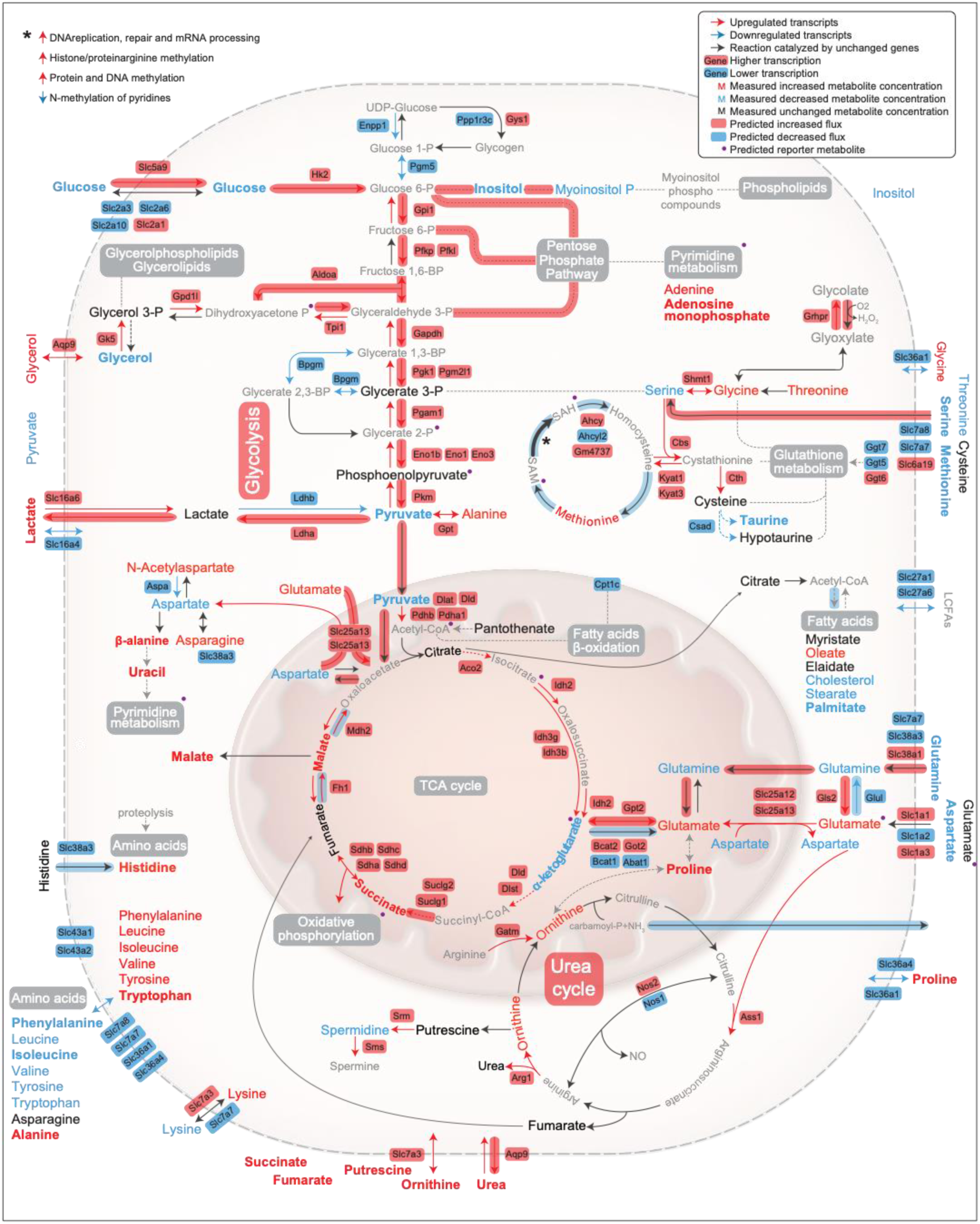
Global overview of altered metabolic pathways in tumor cells, compared to normal in an extended representation. A selection of genes with significantly altered expression (Bonferroni adjusted *p*-value < 0.1; normal n=8, tumor and residual n=4), targeted metabolites with significantly altered levels (Benjamini-Hochberg adjusted *p*-value < 0.01; n=8; WT n=2) and significant reporter metabolites (top 5% with *p*-value < 0.1) of core metabolic processes are presented. Metabolites with an additional log_2_ fold change ≥ 1 or ≤ -1 are highlighted in bold. A Wald test with a Negative Binomial GLM was used as test statistics for gene expression^29^ was used as test statistic for metabolite levels. For the reporter metabolites a gene set enrichment analysis was performed from a theoretical null-distribution using the reporter method^30^. Bonferroni adjusted *p*-values of the expression analysis (Wald test) were used as gene-level statistics. Metabolite-gene sets were derived from a genome wide human metabolic model (HMR2), with genes mapped to mouse orthologs^31^. Flux balance analysis predicted metabolic fluxes, which are altered in the corresponding pathways, are overlaid. Significantly altered gene expressions and metabolites were used to inform the predictions. Number of replicates corresponds to different animals.

**Extended data 7:**
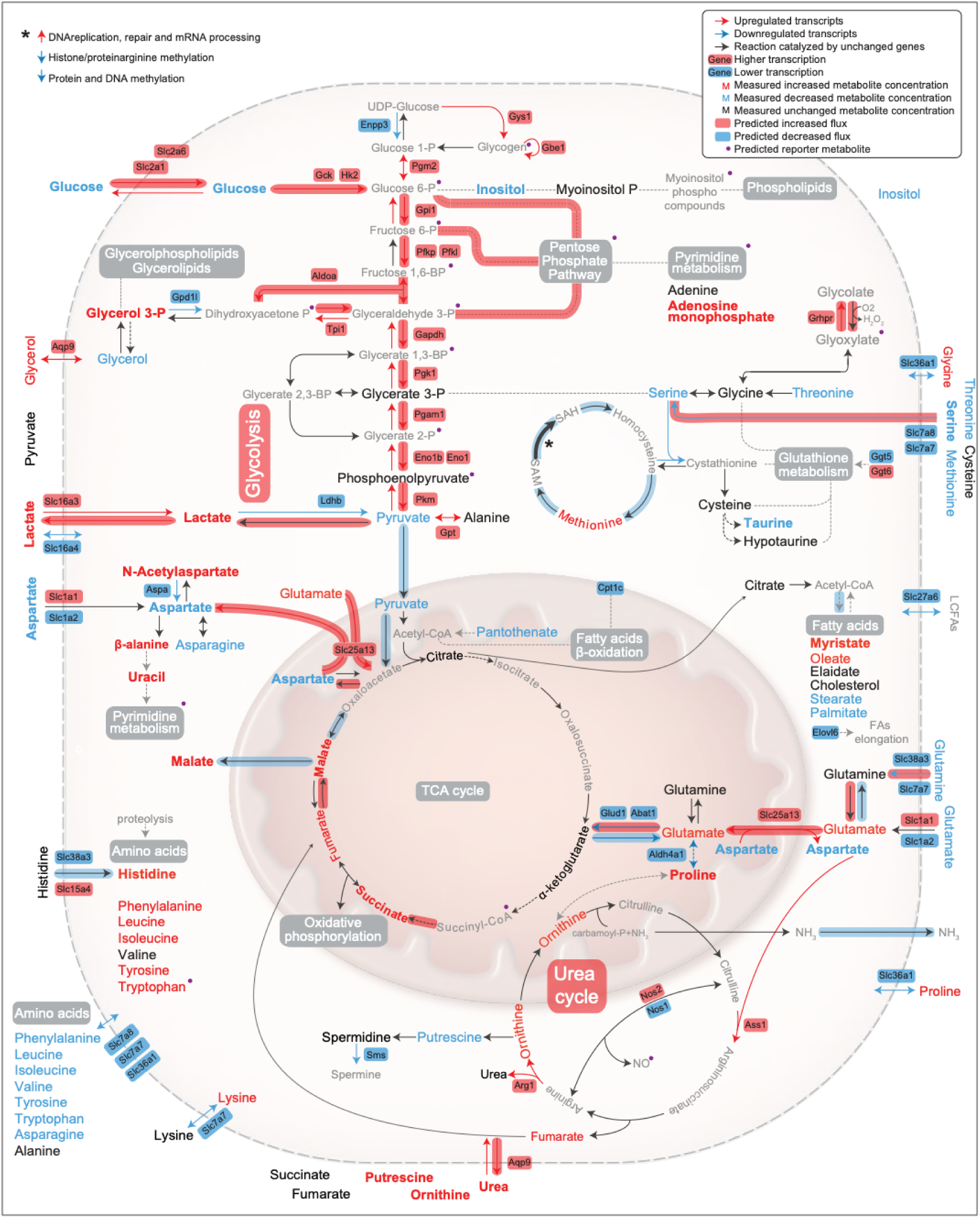
Global overview of altered metabolic pathways in residual cells, compared to normal in an extended representation. A selection of genes with significantly altered expression (Bonferroni adjusted *p*-value < 0.1; normal n=8, tumor and residual n=4), targeted metabolites with significantly altered levels (Benjamini-Hochberg adjusted *p*-value < 0.01; n=8; WT n=2) and significant reporter metabolites (top 5% with *p*-value < 0.1) of core metabolic processes are presented. Metabolites with an additional log_2_ fold change ≥ 1 or ≤ -1 are highlighted in bold. A Wald test with a Negative Binomial GLM was used as test statistics for gene expression^29^ was used as test statistic for metabolite levels. For the reporter metabolites a gene set enrichment analysis was performed from a theoretical null-distribution using the reporter method^30^. Bonferroni adjusted *p*-values of the expression analysis (Wald test) were used as gene-level statistics. Metabolite-gene sets were derived from a genome wide human metabolic model (HMR2), with genes mapped to mouse orthologs^31^. Flux balance analysis predicted metabolic fluxes, which are altered in the corresponding pathways, are overlaid. Significantly altered gene expressions and metabolites were used to inform the predictions. Number of replicates corresponds to different animals.

**Extended data 8:**
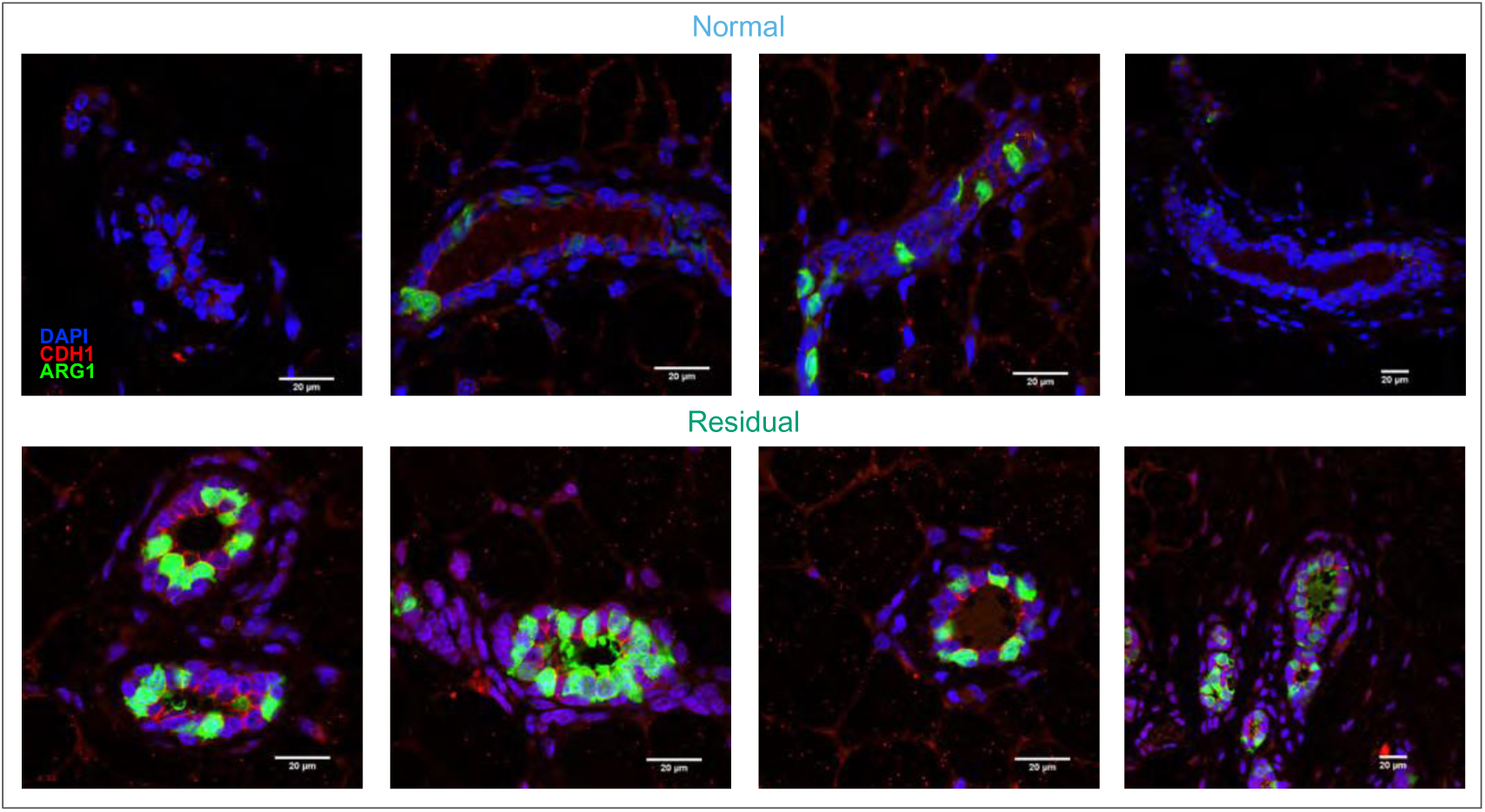
Immunofluorescence staining of ARG1 on tissue sections of healthy (age-matched controls) and regressed mammary glands. Representative images from stainings of healthy (top panel) and residual ducts (panel below). Scale bar: 20 µm Immunofluorescence: DAPI (blue), CDH1 (red), ARG1 (green).

**Supplementary Figure 9:**
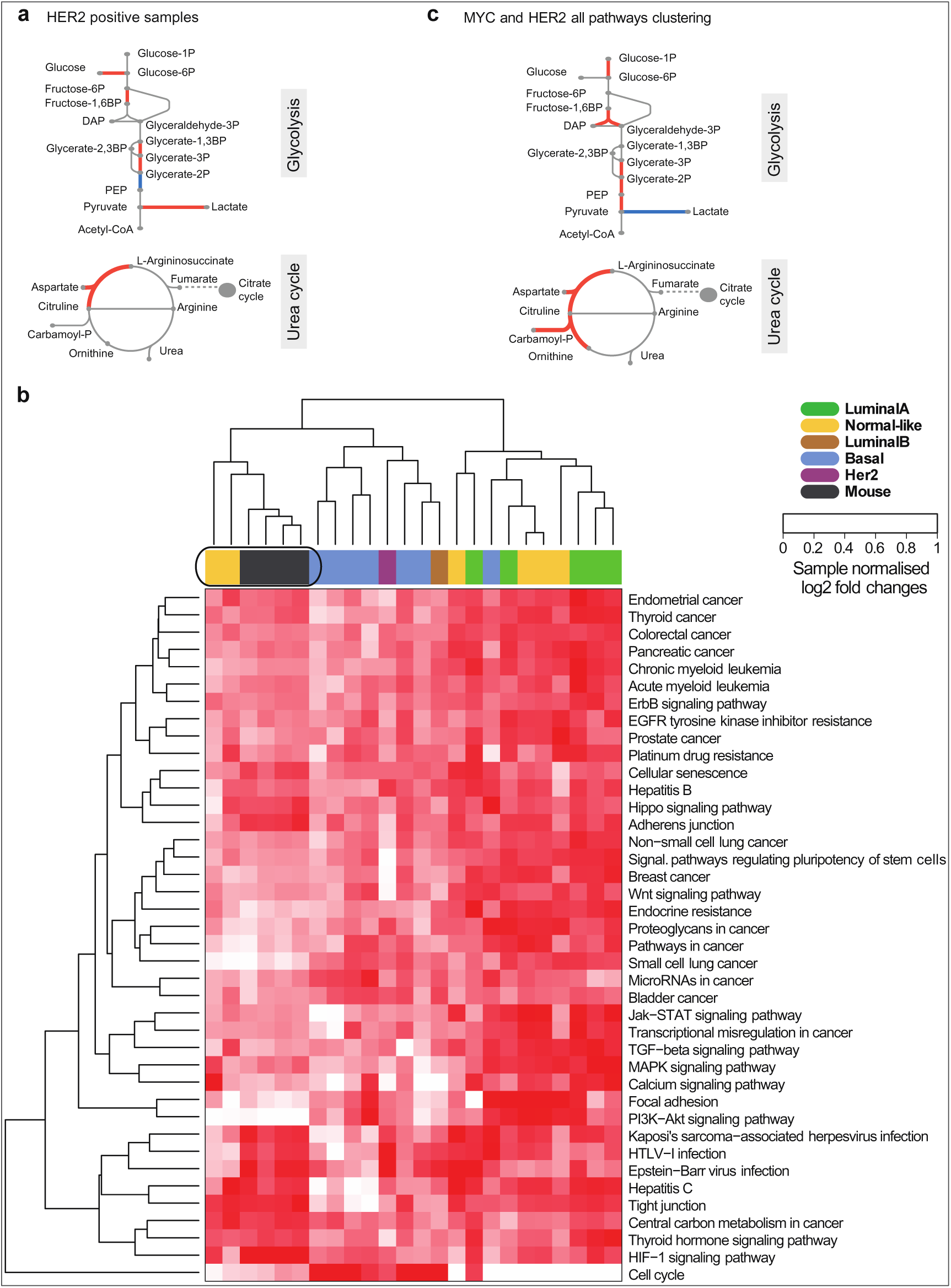
Correlation of mouse data with patient datasets. **a** Metabolic reactions of glycolysis and urea cycle, which are catalyzed by enzymes with differential expression (Benjamini-Hochberg adjusted *p*-value < 0.1), are highlighted in red. An empirical Bayes moderated t-statistics was computed from a gene wise linear model fit with generalized least squares^28^, comparing *HER2* positive classified patients (n=3 samples) with healthy breast tissue (n=10 samples). Differential expression from mouse *in vitro* transcriptome data of residual versus normal samples (RNA-seq, normal n=8, residual n=4) is shown in comparison (Wald test, Bonferroni adjusted *p*-value < 0.1)^29^. **b**, Joint clustering of mouse model (RNA-seq, normal n=8, residual n=4) and patient (microarray, healthy n=10 samples, residual n=20 samples) derived transcriptome log_2_ fold changes of healthy samples compared with regressed tumor samples. The clustering is based on all genes involved in *HER2* or *MYC* related KEGG pathways. Hierarchical clustering with the complete linkage method and the Euclidean distance as a distance metric was used for clustering. For the patient comparison, two independent data sets, one obtained from healthy breast tissue (GSE65194)^32,33^ and one obtained from patient tissues after neoadjuvant treatment (GSE32072)^34^ were merged. **c**, Metabolic reactions of glycolysis and urea cycle, which are catalyzed by enzymes with differential expression (*p*-value < 0.05), are highlighted in red. An empirical Bayes moderated t-statistics was computed from a gene wise linear model fit with generalized least squares^28^, comparing the treated patients (n=4 samples), which are clustering closely with the mouse samples (**b**), with healthy breast tissue (n=10 samples). Differential expression from mouse *in vitro* transcriptome data of residual versus normal samples (RNA-seq, normal n=8, residual n=4) is shown in comparison (Wald test, Bonferroni adjusted *p*-value < 0.1)^29^. **a**-**c**, Number of replicates corresponds to different animals,

**Supplementary Figure 10:**
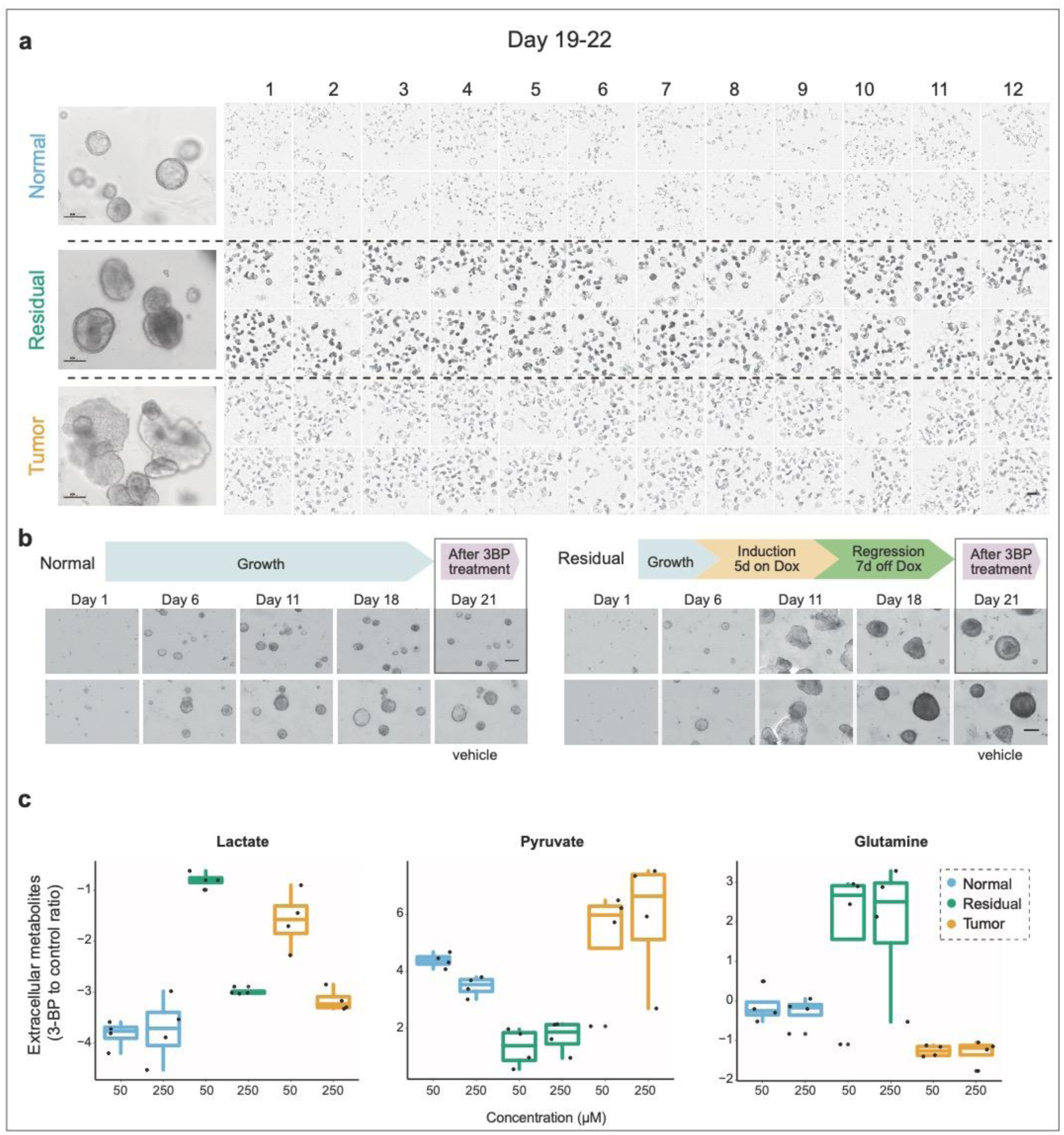
Functional validation of an altered glycolysis in residual cell populations. **a**, Bright field images in transmission showing a 96-well plate containing normal, residual and tumor structures before treatment with 3-BP, scale bar 100 µm (right panel) and 500 µm (96 well plate). **b**, Bright field time-lapse images of the normal (left) and residual (right) structures over the timeline of the experiment: seeding (day 1), growth (day 6), tumorigenesis (day 11), regression (day 18) and after drug treatment (day 21). Scale bar 100 µm **c,** Extracellular metabolite abundance levels alteration in all three populations after treatment with 3-BP (n=4). The values represent the ratio of 3-BP treated to untreated cells. Number of replicates corresponds to different animals.

**Supplementary Table 1.** GO terms for Downregulated genes. Significant gene ontology terms from the gene ontology biological processes for the comparison residual vs normal.

| GO Terms for Down-regulated Genes |  |  |
| --- | --- | --- |
| GO Term ID | Description | Number of significant terms |
| GO:0006260 | DNA replication | 3 |
| GO:0006928 | movement of cell or subcellular component | 1 |
| GO:0007017 | microtubule-based process | 1 |
| GO:0007049 | cell cycle | 1 |
| GO:0007059 | chromosome segregation | 1 |
| GO:0007154 | cell communication | 1 |
| GO:0007155 | cell adhesion | 3 |
| GO:0007188 | adenylate cyclase-modulating G-protein coupled receptor signaling pathway | 2 |
| GO:0007267 | cell-cell signaling | 12 |
| GO:0007610 | behavior | 1 |
| GO:0008283 | cell proliferation | 1 |
| GO:0009190 | cyclic nucleotide biosynthetic process | 16 |
| GO:0009987 | cellular process | 1 |
| GO:0019932 | second-messenger-mediated signaling | 1 |
| GO:0022610 | biological adhesion | 1 |
| GO:0023052 | signaling | 1 |
| GO:0030198 | extracellular matrix organization | 1 |
| GO:0030334 | regulation of cell migration | 8 |
| GO:0032501 | multicellular organismal process | 1 |
| GO:0032502 | developmental process | 1 |
| GO:0042127 | regulation of cell proliferation | 1 |
| GO:0043062 | extracellular structure organization | 1 |
| GO:0044699 | single-organism process | 1 |
| GO:0044707 | single-multicellular organism process | 37 |
| GO:0044708 | single-organism behavior | 1 |
| GO:0044763 | single-organism cellular process | 1 |
| GO:0044767 | single-organism developmental process | 12 |
| GO:0048285 | organelle fission | 17 |
| GO:0048518 | positive regulation of biological process | 1 |
| GO:0048519 | negative regulation of biological process | 3 |
| GO:0048523 | negative regulation of cellular process | 18 |
| GO:0048583 | regulation of response to stimulus | 1 |
| GO:0050896 | response to stimulus | 1 |
| GO:0051301 | cell division | 1 |
| GO:0051302 | regulation of cell division | 1 |
| GO:0051303 | establishment of chromosome localization | 2 |
| GO:0055086 | nucleobase-containing small molecule metabolic process | 1 |
| GO:0065007 | biological regulation | 1 |
| GO:0071467 | cellular response to pH | 1 |
| GO:0071840 | cellular component organization or biogenesis | 1 |
| GO:1903047 | mitotic cell cycle process | 36 |
| GO:1905818 | regulation of chromosome separation | 2 |

**Supplementary Table 2.**
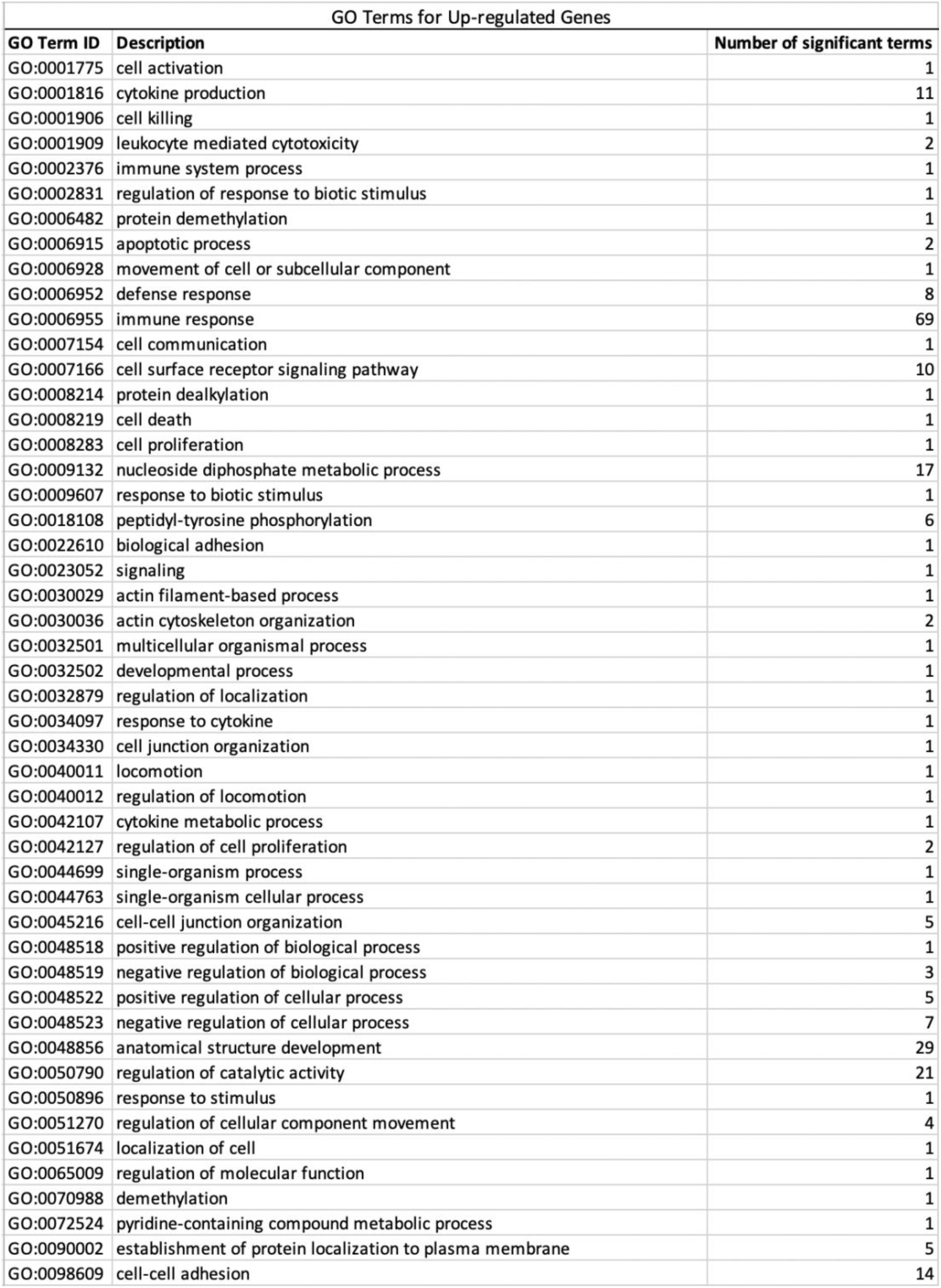
GO terms for Upregulated genes. Significant gene ontology terms from the gene ontology biological processes for the comparison residual vs normal.

| GO Terms for Up-regulated Genes |  |  |
| --- | --- | --- |
| GO Term ID | Description | Number of significant terms |
| GO:0001775 | cell activation | 1 |
| GO:0001816 | cytokine production | 11 |
| GO:0001906 | cell killing | 1 |
| GO:0001909 | leukocyte mediated cytotoxicity | 2 |
| GO:0002376 | immune system process | 1 |
| GO:0002831 | regulation of response to biotic stimulus | 1 |
| GO:0006482 | protein demethylation | 1 |
| GO:0006915 | apoptotic process | 2 |
| GO:0006928 | movement of cell or subcellular component | 1 |
| GO:0006952 | defense response | 8 |
| GO:0006955 | immune response | 69 |
| GO:0007154 | cell communication | 1 |
| GO:0007166 | cell surface receptor signaling pathway | 10 |
| GO:0008214 | protein dealkylation | 1 |
| GO:0008219 | cell death | 1 |
| GO:0008283 | cell proliferation | 1 |
| GO:0009132 | nucleoside diphosphate metabolic process | 17 |
| GO:0009607 | response to biotic stimulus | 1 |
| GO:0018108 | peptidyl-tyrosine phosphorylation | 6 |
| GO:0022610 | biological adhesion | 1 |
| GO:0023052 | signaling | 1 |
| GO:0030029 | actin filament-based process | 1 |
| GO:0030036 | actin cytoskeleton organization | 2 |
| GO:0032501 | multicellular organismal process | 1 |
| GO:0032502 | developmental process | 1 |
| GO:0032879 | regulation of localization | 1 |
| GO:0034097 | response to cytokine | 1 |
| GO:0034330 | cell junction organization | 1 |
| GO:0040011 | locomotion | 1 |
| GO:0040012 | regulation of locomotion | 1 |
| GO:0042107 | cytokine metabolic process | 1 |
| GO:0042127 | regulation of cell proliferation | 2 |
| GO:0044699 | single-organism process | 1 |
| GO:0044763 | single-organism cellular process | 1 |
| GO:0045216 | cell-cell junction organization | 5 |
| GO:0048518 | positive regulation of biological process | 1 |
| GO:0048519 | negative regulation of biological process | 3 |
| GO:0048522 | positive regulation of cellular process | 5 |
| GO:0048523 | negative regulation of cellular process | 7 |
| GO:0048856 | anatomical structure development | 29 |
| GO:0050790 | regulation of catalytic activity | 21 |
| GO:0050896 | response to stimulus | 1 |
| GO:0051270 | regulation of cellular component movement | 4 |
| GO:0051674 | localization of cell | 1 |
| GO:0065009 | regulation of molecular function | 1 |
| GO:0070988 | demethylation | 1 |
| GO:0072524 | pyridine-containing compound metabolic process | 1 |
| GO:0090002 | establishment of protein localization to plasma membrane | 5 |
| GO:0098609 | cell-cell adhesion | 14 |

## Author information

These authors contributed equally: Ksenija Radic Shechter, Eleni Kafkia, Katharina Zirngibl These authors jointly supervised this work: Martin Jechlinger, Kiran Raosaheb Pati

## Contributions

K.R.S. and A.A. carried out mouse work, optimized the 3D culture system and characterized it through immunofluorescent staining and RNA-sequencing. K.R.S. and E.K. optimized and performed all metabolic experiments. K.Z. provided statistics and analysis of multi-omics data and mouse-human transcriptome data comparison. D.M. and K.Z. performed genome-scale metabolic modeling and flux balance analysis. D.C.S. performed untargeted metabolomics analysis. E.K optimized and analyzed targeted metabolomics and ^13^C tracer experiments. C.L. and B.B. performed lipidomics analyses. K.R.S. and E.K. carried out NOS enzymatic assay. K.R.S. performed immunohistochemical and immunofluorescence staining on mouse tissue sections. K.R.S. and S.G. designed and performed glycolysis inhibition experiments. S.G. analyzed CellTox and ScanR data and contributed to manuscript preparation. K.R.S. collected and extracted DNA for methylation experiments. K.Z. analyzed DNA methylation data. K.R.S., E.K. and K.Z. carried out study design, interpretation and manuscript preparation. M.J. and K.R.P. provided study design, supervision, interpretation, manuscript preparation and critical review.

## Competing interests

The authors declare no competing interests.

## Acknowledgements

The authors want to thank Marta Garcia Montero, Savannah Jackson, Matthew Boucher and Dr. Lucas Chaible for their help in harvesting material for *in vivo* experiments, technical assistance with histology and maintenance of the mouse colony; Federico Villa for contribution to metabolic methods setup; Dr. Yuanyuan Chen for the help in analysis of in *vivo* fluorescence staining data at the Tissue-gnostic microscope and the group of Dr. Rocio Sotillo at DKFZ Heidelberg for technical support; Dr. Vladimir Benes and Ferris Jung from the Genomic Core Facility at EMBL for their expert advice and technical assistance on RNA sequencing and bisulfite-free DNA sequencing; Kerstin Putzker, Eugene Gbekor and Dr. Joe Lewis from Chemical Core Facility for expertise with the inhibitor assays design and technical assistance; Sabine Reither, Stefan Terjung from Advanced Light Microscopy Facility at EMBLfor assistance with the image acquisition; Christian Tischer from Center of Bioimage Analysis at EMBL; Laboratory Animal Resources for help with mouse work and colony maintenance; Johanna Vappiani for untargeted metabolomics mass spectrometry; Arnaud Krebs for help with the methylation analysis; Bernd Klaus for general statistical analyses help; Sergej Andrejev for help with the untargeted metabolomics analysis. S.G. was supported by a fellowship from the EMBL Interdisciplinary Postdoc (EI3POD) programme under Marie Sklodowska Curie Actions COFUND (grant agreement number 664726); Metabolomics analysis received funding through EMBL-Merck Serono GmbH (Darmstadt, Germany) collaboration program; B.B. was funded by the Deutsche Forschungsgemeinschaft (DFG, German Research Foundation) - Project Number 112927078 - TRR 83 and Project Number - 331351713 - SFB1324; M.J. was funded by Marie Curie PCIG-GA-2011-294121, 3DBreastCancer.

## Materials and Methods

### Animals

Breeding and maintenance of mouse colony was done in LAR (Laboratory Animal Resources) facility of EMBL Heidelberg, in accordance to the guidelines of the European Commission, revised Directive 2010/63/EU and AVMA Guidelines 2007, under veterinarian supervision. Animals – *TetO-cMYC/TetO-Neu/MMTV-rtTA*^1,2^ in FVB background – were kept on a 12-hour light/12-hour dark cycle, with constant ambient temperature (23±1°C) and humidity (60±8%), supplied with food pellets (for tumor induction, pellets contained doxycycline hyclate, 625 mg/kg; Envigo Teklad) and water *ad libitum*.

For purpose of genotyping, genomic DNA was extracted by tail-digestion in 75 µl of digestion buffer (NaOH 25 mM + EDTA 0,2 mM) at 98°C followed by addition of 75 µl Tris-HCl (40 mM, pH 5,5) and centrifugation at 4000 rpm for 3 min. Gel-electrophoresis was used for the detection of PCR products (*MYC* 630 bp, Neu 386 bp, rtTA 380 bp) on 1,5% agarose (Sigma, A9539-500G) gel with Ethidium bromide solution (Sigma, E1510-10ML) in a final concentration of 0,5 µg/ml The products were visualized using Quantum-Capt1 documentation system (Peqlab).

### 3D cell cultures

Three-dimensional cell cultures were established according to the published protocol^3^ with some modifications. Primary mammary epithelial cells were obtained from 8 weeks old virgin females of described mouse strains, through digestion of mammary glands in 5 mL of digestion media (Lonza/Amaxa DMEM/F12 1:1 Mixture with HEPES, L-Gln, BE12-719F), supplemented with HEPES to the final concentration of 25 mM, 150 U/mL Collagenase type 3 (Worthington, LS004183), 20 µg/mL Liberase Blendzyme 2 (Roche, 05401020001) and 5 ml of Penicillin/Streptomycin (Gibco Life Technologies, 15140-122). Digestion for 15-16 hours at 37°C in 5% (vol/vol) CO_2_ atmopshere, in loosely capped 50 mL polypropylene conical tubes, was followed by washing step with 45 mL of phosphate-buffered saline (PBS). Upon centrifugation at room temperature, 1000 rpm for 5 min, interphase between upper fat layer and cell pellet was removed and 5 mL of 0.25% trypsin-EDTA (Invitrogen, 25200-056) was added. Suspension was incubated for 40 min at 37°C, 5% CO_2_ in loosely capped tubes, followed by the wash with 25 mL of STOP media (Lonza/Amaxa DMEM/F12 1:1 Mixture with HEPES, L-Gln, BE12-719F supplemented with HEPES to the final concentration of 25 mM and 10% Tet System Approved Fetal Bovine Serum, Biowest, S181T) and treatment with 5-15 mg/mL DNase I (ThermoFisher, 18068015). After another centrifugation step at room temperature, 1000 rpm for 5 min, dissociated cells were resuspended in MEBM media (Lonza, Mammary Epithelial Cell Basal Medium CC-3151 with supplements from Mammary Epithelial Cell Medium BulletKit CC-3150) and plated onto collagen-coated plates (BD Biosciences, 356400) for selection of epithelial cells. Next day cells were washed with PBS and the remaining ones treated with 500 µl of 0,25 % trypsin-EDTA until detachment. Trypsin was inactivated with 9 mL of STOP media (described above), followed by centrifugation step at room temperature, 1000 rpm for 5 min. Cell pellets were resuspended in PBS, counted, and mixed rapidly on ice with the prepared Matrigel-collagen mixture – Cultrex 3D Culture Matrix Basement Membrane Extract (Biozol, TRE-3445-005-01) and 1,5 mg/mL Cultrex 3D Collagen I rat tail (TEMA Ricerca, 3447-020-01). Mixed droplets in volume of 100 µl, containing 12 000 primary mouse mammary epithelial cells, were dispensed into flat bottom wells (Corning CellBIND 12 Well Clear Multiple Well Plates, 3336) or chambered cover glass slides for imaging (ThermoFisher Scientific, Nunc LabTek II Chambered Cover glass, 155379). After solidifying for 40 min at 37°C, 1.5 ml of MEBM serum-free media (supplemented with 2 mL of bovine pituitary extract, 0.5 mL of hEGF, 0.5 mL of hydrocortisone, 0.5 mL of GA-1000, 0.5 mL Insulin from Mammary Epithelial Cell Medium BulletKit CC-3150) was added to each well. Doxycycline (Sigma, Doxycycline hyclate, D9891) was added in concentration 200 ng/ml. For experiments based on biochemical assays, suspension of 500 cells and PBS was mixed with Matrigel in ratio 1:4 for 5 µl gels, that were seeded in the 96-well Corning black polystyrene microplates (Sigma-Aldrich, CLS3603) and left to solidify for 15 min at 37°C, followed by addition of 100 µl of MEBM media. Reseeding experiments were done by incubation of 100 µl gels with collagenase and liberase for 1,5 h at 37°C which led to digestion of Matrigel, releasing the structures from the gel. After washing with PBS, cells were incubated for 5-10 min at 37°C with Trypsin (150 µl per gel), washed with previously described STOP media, counted and seeded in microplates as 5 µl gels.

### Immunofluorescence

3D culture gels for immunofluorescence staining were fixed with 4% paraformaldehyde (PFA) for 7-10 min and transferred to the IF deactivated clear glass screw neck vials (Waters, 186000989DV), washed three times with PBS and once in IF buffer (containing NaCl, Na_2_HPO_4_, NaN_3_, BSA, TritonX-100, Tween-20; pH 7,4). Blocking was done using 1x IF buffer with 10% goat serum (Jackson Immuno Research, 005-000-121) for 1,5 h, followed by incubation at 4°C overnight. Primary antibodies were diluted in primary block as described above and washed the next day in 1x IF buffer 3x, 15 min each. Incubation with secondary antibodies and 4’, 6’-diamino-2-phenylindole (DAPI) was done in the primary block for 1 hour (dilution 1:1000). Gels were washed with 1x IF buffer and 1x PBS, 2 times, 10 min each. Gels were mounted with Vectashield Anti-fade mounting medium (Vinci Biochem, VC-H-1500-L010) into LabTek II chamber slide (ThermoFisher Scientific, 50733) and imaged on a Leica SP5 confocal microscope using 63x water lens and LAS AF imaging software. The following antibodies were used for the 3D cultures: alpha-6-integrin (BD Biosciences 25-0495-82, diluted 1:80), ZO-1 (Invitrogen 61-7300, diluted 1:500), GM-130 (BD Biosciences, 610823, diluted 1:100), E-cadherin (Invitrogen, 13-1900, diluted 1:200). Nuclei were stained with DAPI (ThermoScientific, 62248, diluted 1:1000). Anti-rabbit, anti-mouse, and anti-rat antibodies were purchased coupled with Alexa Fluor dyes from Invitrogen (A21247, A11034, A11036). FFPE tissue sections were stained using the standard protocols for ARG1 (Novus, NBP1-32731, diluted 1:250) antibody. Sections were mounted using ProLong Gold Antifade Mountant (ThermoFisher Scientific, P36930) and scanned using TissueFAXS Slides system (TissueGnostics). Quantification was done using StrataQuest Analysis Software (TissueGnostics).

### Immunohistochemistry

MT-CO1 antibody (Abcam, ab45918) staining was done on FFPE tissue sections following the standard IHC protocol: deparaffinization and rehydration of the samples; antigen retrieval using citric acid-based Antigen unmasking solution (Vector, H-3300) for 30 min in a steamer and inactivation of endogenous hydrogen peroxidase activity with 10% H_2_O_2_ solution (Sigma, H1009), followed by blocking using 10% Normal goat serum (Jackson Immuno Research, 00500121) in 1x Phosphate Saline Buffer (PBS).

Incubation with primary antibody (MT-CO1 diluted 1:500) was done in blocking buffer at 4°C overnight, after which the slides were washed 3x, 5 min each, using 1x PBS, followed by incubation with biotinylated antibody (Peroxidase, Rabbit IgG; Vector Laboratories, PK-6101) for 30 min, washing and incubation with Horse Radish Peroxidase (HRP) conjugated antibody and detection using DAB Peroxidase (HRP) Substrate Kit (Vector, SK-4100). Counter-staining was done using Hematoxylin QS (Vector, H-3404), after which the sections were dehydrated, mounted with DPX Mountant for histology (Sigma, 06522) and analyzed using LMD 7000 microscope (Leica) equipped with Leica CD310 digital camera and LASV3.7 (Leica) software.

### RNA collection and extraction

RNA was harvested from a pool of two 3D gels per condition, using 900 µl of mirVana lysis buffer. Extraction was done using mirVana miRNA Isolation Kit, with phenol (Ambion, AM1560). After assessing RNA quality and concentration on Bioanalyzer (Agilent 2100, G2939BA), RNA was sequenced in the Genomics Core Facility at EMBL on Illumina NextSeq 500 platform, non-directional single end read length NextSeqHigh 75 bp.

### Analysis of RNA sequencing data

After assessing the quality of the raw RNA sequencing reads by FastQC version 0.11.3. [Andrews S. FastQC: a quality control tool for high throughput sequence data. 2010. Available at: http://www.bioinformatics.babraham.ac.uk/projects/fastqc/], adapter trimming using Cutadapt version 1.9.1 [Martin M. Cutadapt removes adapter sequences from high-throughput sequencing reads. EMBnet.journal. 2011;17(1):10–2.] with default options providing the standard Illumina TrueSeq Index adapters was done. FaQCs version 1.34^4^ was used for subsequent quality trimming and filtering, applying the following parameters: -q 20 -min_L 30 -n 5 -discard 1. Total reads per sample after trimming and filtering ranged from 34.1 to 52.0 million. Sequencing reads were aligned to the M. musculus reference genome (GRCm38.p4) [NCBI. Genome Reference Consortium Mouse Build 38 patch release 4 (GRCm38.p4) Available at: https://www.ncbi.nlm.nih.gov/assembly/GCF_000001635.24/] using Tophat2 version 2.0.10^5^, which included the sequence for human c-MYC and rat HER2 with the following parameter: -G -T -x 20 -M --microexon-search --no-coverage-search --no-novel-juncs -- mate-std-dev 100 -r 50 --min-segment-intron 20 -i 30 –a 6. For the differential expression analysis, only reads with unique mappings were considered. Gene level count tables were obtained using the count script of the HTSeq python library version 0.6.1p1 ^6^ with default options. All reads mapped in total to 19500 to 20800 genes across all samples. For performing a dimensionality reduction by principal component analysis (PCA) and hierarchical clustering rlog transcript counts were utilized, transformed with the “rlog” function of the Bioconductor package DESeq2 version 1.12.4 (10). R V.3.3.1. (R Development Core Team) was used for conducting biostatistical analyses.

### Differential expression analysis

The statistical analysis for differential expression was mainly done with the Bioconductor package DESeq2 version 1.12.4 ^7^. Size-factor based normalization was performed to control for batch effects and inter-sample variability. Genes with less than 10 counts across all samples were filtered to increase the sensitivity of the detection of differential gene expression. Package defaults were used for dispersion estimation and differential expression analysis with the function “Deseq”, which includes independent filtering, cooks cutoff^8^ for outlier detection and the performance of a Wald-test. The animal was included as a confounder variable in the model design. Subsequently, adjusted p-values were computed from the DESeq2 calculated p-values by applying a Bonferroni correction for multiple testing. Genes with a p_adj_-value < 0.1 were considered as significantly differentially expressed (DE). R V.3.3.1. (R Development Core Team) was used for conducting biostatistical analyses.

### Gene set enrichment analyses

2039 differentially expressed genes in residual (compared to never induced control; q-value < 0.1) and 6411 differentially expressed genes in tumor cells (compared to never induced, q-value < 0.1) were taken for gene ontology (GO) enrichment analysis. GO enrichment analysis was performed using Fisher’s exact test with a foreground of all respective differentially expressed genes and a background, which was composed of a unique set of 5 randomly picked genes per foreground gene exhibiting a similar expression mean over all samples. The analysis was done separately for up-regulated and down-regulated genes. The chosen cutoff for significant GO terms was p-value < 0.001. Further, a gene set enrichment analysis for significantly enriched KEGG pathways was performed for up-regulated and down-regulated differentially expressed genes separately using the function “gage” of the likewise called R package with version 2.32.1 ^9^. The function calculates sample-wise test statistics with an unpaired two-sample t-test using annotations from “org.Mm.eg.db” version 3.7.0 [Marc Carlson (2018). org.Mm.eg.db: Genome wide annotation for Mouse. R package version 3.7.0.]. KEGG pathways with a p-value < 0.05 were considered significantly enriched.

Additionally, a reporter metabolite analysis was performed to identify metabolites or metabolic pathways that are likely to be de-regulated. Therefore, the q-values and log2 fold changes (FC) of the respective differentially expressed genes were used to calculate p-values from a theoretical null distribution (10000 permutations) utilizing the reporter metabolite algorithm from the “piano” R package^10^. Multiple testing adjustment was applied using the Benjamini-Hochberg procedure. The threshold for significance was p_adj_-value < 0.01 for the non-directional class, the distinct-directional class and the mixed-directional class, but maximally the top 5% of the total list of tested metabolites considering each class equally. Pathway enrichment was calculated for gene sets of 1 gene per group or bigger. The gene set was obtained from a revised version of the HMR2 model^11^, whose gene-protein-reaction annotations were translated to mouse orthologs. R V.3.3.1. (R Development Core Team) was used for conducting biostatistical analyses.

### Integration of transcriptomics data into flux balance analysis

Differential gene expression data was integrated info flux balance analysis (FBA) using a new simulation method, Metabolic Analysis with Relative Gene Expression (MARGE). This method aims to overcome some limitations identified in other previously published methods^12^. In particular, it avoids making assumptions on any direct proportionality between transcript levels and reaction rates, instead it uses relative expression between two conditions, as an indication of the direction and magnitude of the flux control exerted on a metabolic pathway through transcriptional regulation. The implementation is based on a previously-proposed extension of FBA that integrates gene-protein-reaction (GPR) association rules into the stoichiometric matrix of the metabolic network, allowing the computation of enzyme-specific flux rates^13^, and is formulated as two-step linear optimization problem. The first step optimizes the agreement between relative enzyme usage and relative gene expression, and the second adds a parsimonious enzyme usage criterion.

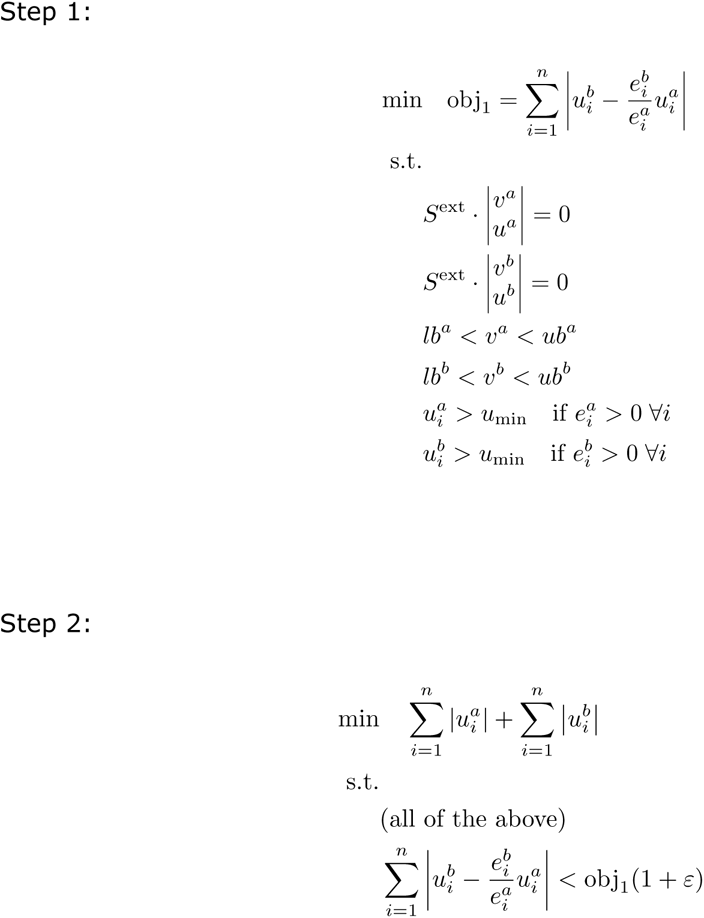

Where *a* and *b* are two experimental conditions, *e^a^* and *e^b^* are the gene expression in each condition, *v^a^* and *v^b^* are reaction flux vectors, *u^a^* and *u^b^* are enzyme usage vectors, *S^ext^* is the extended stoichiometric matrix, *u_min_* is a flux activation threshold (set to 0.001), and *ɛ* is a relaxation term with regard to the first objective (set to 0.1).

### Genome-scale metabolic modeling

Transcriptomics and extracellular GCMS metabolomics data from this study were used as model inputs to refine the phenotype predictions performed with flux balance analysis. The transcriptomics data were provided as log2 fold changes of significantly differentially expressed metabolic genes (q-value < 0.1) between the two experimental conditions. The metabolomics data were used to constrain the metabolite uptake/secretion rates in the model, both in terms of absolute rates per condition, and relative rates between conditions. The fold change of significantly changed extracellular metabolite profiles (*p*_adj_-value < 0.01) were calculated and imposed as constraints in the model with a deviation tolerance of 50% to account for measurement error. For metabolites whose level did not significantly change between conditions, absolute constraints were imposed to ensure a minimum level of uptake/secretion in accordance with the observed directionality. Measurements from pure media allowed determining active secretion/uptake of metabolites into/from the media in respect to both conditions.

A revised version of the human genome-scale metabolic model HMR2 (14) was used for simulation, in which mouse gene orthologs of the gene-protein-reaction annotations had been updated/corrected as well as the model itself to yield more accurate flux predictions. In brief, this included the introduction of a mitochondrial intra-membrane space (adapted from^14^) to improve the prediction of respiratory ATP synthesis, the revision of reactions from the beta-oxidation pathway and auxiliary enzymes, the introduction of ATP maintenance costs and the adaption of model uptakes and releases of metabolites from experimental data^15^. Further, atomically unbalanced reactions were removed and the directionality of reactions was constraint where infeasible. The import/export of SBML files was obtained through the libSBML API^16^ using the load_cbmodel of reframed. The IBM ILOG CPLEX Optimizer version 12.8.0 was used for solving the MILP problems. All simulations were conducted with Python 3.6.9.

The simulations were performed using the *reframed* python package (version 1.0.0) (https://doi.org/10.5281/zenodo.3478380). In particular, we used the MARGE function (implementing the method described above) with the following parameter settings: growth_frac_a=0.8, growth_frac_b=0.8, activation_frac=0.001, step2_tol=0.1. The IBM ILOG CPLEX Optimizer (version 12.8.0) was used for solving the MILP problems. All simulations were conducted with Python 3.6.9.

### Human breast cancer transciptome comparison

Microarray gene expression datasets from breast cancer patients pre- and post-treatment^17^ and control breast tissue from healthy women^18^ were downloaded from Gene Expression Omnibus (GEO)^19^. Each dataset was first analyzed independently, which included filtering for sample outliers, normalization, background correction, minimal intensity filtering of genes and the annotation of genes from probe set IDs with the removal of multiple mappings of transcript cluster identifiers. The sample outliers were identified with the function “arrayQualityMetrics” of the likewise called R package with version 3.38.0^20^. Normalization and background correction were done using the “rma” function of the R package “oligo” version 1.46.0^21^. The minimal intensity threshold for gene filtering was determined by fitting a null model to the whole data set and taking the lower 5% boarder as a cut off. The two datasets were then combined and processed together a second time (normalization, outlier removal, intensity filtering). In order to address the batch effect of the joined data stemming from the two experimental settings, the first principal component was removed from the data set. In addition, the “normal-like” tumor subtype of the patients’ dataset was removed due to the poorly defined diagnostic category and high biological variability. For the differential gene expression a gene-wise linear model was fitted to the dataset using generalized least squares and including the tumor subtype as a confounder variable if applicable. Next an empirical Bayes moderated t-statistic and log-odds were computed with the “eBayes” function of the “limma” package version 3.38.3^22,23^ using the package defaults. All genes with a Benjamini-Hochberg adjusted *p*-value < 0.1 were considered differentially expressed. R V.3.3.1. (R Development Core Team) was used for conducting biostatistical analyses.

### Intracellular and extracellular sample harvest and metabolite extraction

#### 3D cell cultures

Prior to the harvest of organoids, 50 µl of spent growth media were snap-frozen and stored at -80°C until the extraction of metabolites for extracellular metabolomics. Following the collection of the spent growth media, the organoid structures were freed from Matrigel upon digestion for 1,5h- at 37°C with liberase and collagenase added to the remaining media. Subsequently, the media were aspired, the organoids were washed three times with PBS, centrifuged shortly (1000 r.p.m, 2 min, at room temperature) and quenched with 200 µl cold (−80°C) HPLC-grade methanol (Biosolve Chimie, 136841). For metabolite extraction, adonitol (Alfa Aesar, 488-81-3) was added as an internal standard to the organoids/methanol mixture and the samples were incubated at 72°C for 15 min, followed by addition of 200 µl ice-old MilliQ water and centrifugation (15000 r.p.m, 10 min, 4°C). The supernatants were collected and dried with a speed-vac. The dried metabolite extracts were stored at -80°C until metabolomics analysis. Metabolite extraction from the spent growth media was performed as described above by adjusting the volume of the extraction solvents to 100 µl of HPLC-grade methanol and 100 µl of MilliQ water. Finally, 50 µl of the initial pure growth media as well as from the last washing solution were collected and extracted as described above, with the latter sample employed as control to validate the effective washing of the organoids from the extracellular media before quenching.

#### In vivo and ex vivo mammary glands experiments

For experiments that allowed for tumorigenesis and regression in vivo, food pellets supplemented with doxycycline (625 mg/kg) were used for tumor induction in mice which were weekly monitored for tumor detection and their overall health. Full blown tumors developed in the period of 4-6 weeks and when burden was too large (d = 2 cm), animals were given food without doxycycline which resulted in the fast tumor regression to a non-palpable state. At the timepoint of the complete tumor regression (9 weeks after oncogenes deactivation), mammary glands were harvested from these mice, along with the wild-type (non-inducible) siblings which had the same treatment. Before harvesting, vaginal lavage to check for the phase of estrous cycle was obtained, according to the modified protocol^24^: slides were dried at room temperature, fixed in 10 % formalin, washed in 1x PBS, stained with Crystal Violet solution (Sigma, V5265) followed by wash in water and analyzed using Leica Application Suite X and Leica DFC7000 T microscope (Leica Microsystems). For [U-^13^C]glucose tracing experiments, mammary glands were dissected, minced and digested for 2 hours at 37°C using collagenase and liberase enzymes, then cultured for 8 hours at 37°C in 5% (vol/vol) CO_2_ atmopshere, in DMEM glucose- and pyruvate-free media (ThermoFisher Scientific, 11966025) supplemented with 4,5 g/L labeled D-Glucose U-13C, 99% (Cambridge Isotope Laboratories, Inc., CLM-1396-1) and 2 mL of bovine pituitary extract, 0.5 mL of hEGF, 0.5 mL of hydrocortisone, 0.5 mL of GA-1000, 0.5 mL insulin from Mammary Epithelial Cell Medium BulletKit CC-3150. For non-labeled metabolomics experiment, mammary glands were dissected and cultured for 8 hours at 37°C in 5% (vol/vol) CO_2_ atmosphere, in DMEM, High Glucose (4,5 g/L glucose) GlutaMAX (Gibco, 10569044) supplemented with 2 mL of bovine pituitary extract, 0.5 mL of hEGF, 0.5 mL of hydrocortisone, 0.5 mL of GA-1000, 0.5 mL insulin from Mammary Epithelial Cell Medium BulletKit CC-3150. The spent growth media were collected and extracted for metabolomics analyses as described above. For intracellular metabolomics sample harvesting, the mammary glands were collected following the cultivation period, washed two times with PBS and quenched with 200 µl of cold (−80°C) HPLC-grade methanol. Subsequently, the metabolite extraction was performed as described for the 3D cell cultures.

#### Targeted metabolomics analysis with Gas chromatography – Mass Spectrometry

Dried metabolite extracts were derivatized with 50 µl of 20 mg/mL methoxyamine hydrochloride (Alfa Aesar, 593-56-6) solution in pyridine (SigmaAldrich, 437611) for 90 min at 40°C, followed by addition of 100 µl N-methyl-trimethylsilyl-trifluoroacetamide (MSTFA) (Alfa Aesar, 24589-78-4) for 12 hours at room temperature^25,26^. GC-MS analysis was performed using a Shimadzu TQ8040 GC-(triple quadrupole) MS system (Shimadzu Corp.) equipped with a 30m x 0.25 mm x 0.25 µm DB-50MS capillary column (Phenomenex, USA). 1 µl of sample was injected in split mode (split ratio 1:10) at 250^0^C using helium as a carrier gas with a flow rate of 1 ml/min. GC oven temperature was held at 100^0^C for 4 min followed by an increase to 320^0^C with a rate of 10^0^C/min, and a final constant temperature period at 320^0^C for 11 min. The interface and the ion source were held at 280^0^C and 230^0^C, respectively. The detector was operated both in scanning mode recording in the range of 50-600 m/z, as well as in MRM mode for specified metabolites. The metabolite identification was based on an in-house database with analytical standards being utilized to define the retention time and the mass spectrum for all the quantified metabolites. The metabolite quantification was carried out by calculating the area under the curve (AUC) of the marker ion of each metabolite normalized to the AUC of adonitol’s marker ion 319. Subsequently, the AUCs were normalized to total metabolite levels. To identify the statistically significant altered metabolites the limma package^22^ (version 3.36.5) in R (version 3.5.2) was utilized with the significance threshold corresponding to a Benjamini-Hochberg adjusted p value ≤ 0.01. Metabolites that followed the additional criterion of a log_2_ fold change (regressed or tumor compared to healthy) ≥ 1 or ≤ -1 were further highlighted in the plots with bold letters.

For the [U-^13^C]glucose tracing experiments, dried metabolite extracts were derivatized with 50 µl of 20 mg/mL methoxyamine hydrochloride (Alfa Aesar, 593-56-6) solution in pyridine (SigmaAldrich, 437611) for 90 min at 40°C, followed by addition of 100 µl N-tert-Butyldimethylsilyl-N-methyltrifluoroacetamide + 1% tert-Butyldimethylchlorosilane (Sigma Aldrich, 00942) for 1 hour at 60°C. The samples remained at room temperature until GC-MS analysis. The GC-MS was operated using the same conditions as described above with the following difference: GC oven temperature was held at 100^0^C for 3 min followed by an increase to 300^0^C with a rate of 3.5^0^C/min, and a final constant temperature period at 300^0^C for 10 min. The natural abundance isotopes were corrected using the Isotope Correction Toolbox (ICT)^27^. Statistics were calculated by unpaired two-samples t-test following assessment of normality and equal variance using the Shapiro-Wilk’s test and F test, respectively.

#### Untargeted metabolomics by flow injection mass spectrometry

Untargeted metabolomics analysis was performed based on a previously published approach^28^. Briefly, samples were analyzed on a LC-MS platform consisting of a Thermo Scientific Ultimate 3000 liquid chromatography system with autosampler temperature set to 10° C coupled to a Thermo Scientific Q-Exactive Plus mass spectrometer equipped with a heated electrospray ion source and operated in negative ionization mode. The isocratic flow rate was 150 µL/min of mobile phase consisting of 60:40% (v/v) isopropanol:water buffered with 1 mM ammonium fluoride at pH 9 and containing 10 nM taurocholic acid and 20 nM homotaurine as lock masses. Mass spectra were recorded in profile mode from 50 to 1,000 m/z with the following instrument settings: sheath gas, 35 a.u.; aux gas, 10 a.u.; aux gas heater, 200° C; sweep gas, 1 a.u.; spray voltage, -3 kV; capillary temperature, 250° C; S-lens RF level, 50 a.u; resolution, 70k @ 200 m/z; AGC target, 3×10^6^ ions, max. inject time, 120 ms; acquisition duration, 60 s. Spectral data processing was performed using an automated pipeline in R. Detected ions were tentatively annotated as metabolites based on matching accurate masses of assumed [M-H] and [M-2H] ions with either no, one or two ^12^C to ^13^C exchanges within a tolerance of 5 mDa to compounds in the Human Metabolome database as reference^29^, with the method-inherent limitation of being unable to distinguish between isomers. Hierarchical clustering of the ions was performed with the complete linkage method and the Euclidean distance as a distance metric. Only ions with non-zero intensity and unique annotation including no assumed mass shift were used for the clustering analysis. For the visualization, ions with a false discovery rate < 0.05 as determine by unpaired two-sided t-tests and subsequent multiple hypothesis testing correction according to Storey’s and Tibshirani’s method^30^ were considered as significantly changed.

#### Lipidomics

Acidic extractions were performed as described^31^ in the presence of an internal lipid standard mix containing 50 pmol phosphatidylcholine (13:0/13:0, 14:0/14:0, 20:0/20:0; 21:0/21:0, Avanti Polar Lipids), 50 pmol sphingomyelin (d18:1 with N-acylated 13:0, 17:0, 25:0), 100 pmol D6-cholesterol (Cambridge Isotope Laboratory), 25 pmol phosphatidylinositol (16:0/ 16:0, Avanti Polar Lipids), 25 pmol phosphatidylethanolamine and 25 pmol phosphatidylserine (both 14:1/14:1, 20:1/20:1, 22:1/22:1), 25 pmol diacylglycerol (17:0/17:0, Larodan), 25 pmol cholesteryl ester (9:0, 19:0, Sigma), 24 pmol triacylglycerol (D5-Mix, LM-6000/D5-17:0/17:1/17:1, Avanti Polar Lipids), 5 pmol ceramide and 5 pmol glucosylceramide (both d18:1 with N-acylated 15:0, 17:0, 25:0), 5 pmol lactosylceramide (d18:1 with N-acylated C17 fatty acid, Avanti Polar Lipids), 10 pmol phosphatidic acid (21:0/22:6, Avanti Polar Lipids), 10 pmol phosphatidylglycerol (14:1/14:1, 20:1/20:1, 22:1/22:1), 10 pmol lyso-phosphatidylcholine (17:1, Avanti Polar Lipids), 50 pmol cardiolipin (14:1/14:1/14:1/15:1, Avanti Polar Lipids), and 50 pmol monolysocardiolipin (16:0/16:0/16:0, Avanti Polar Lipids). Neutral extractions were performed as described^31^ containing a phosphatidylethanolamine plasmalogen (PE P-)-standard mix which was spiked with 16.5 pmol PE P-Mix 1 (16:0p/15:0, 16:0p/19:0, 16:0p/ 25:0), 23.25 pmol PE P-Mix 2 (18:0p/15:0, 18:0p/19:0, 18:0p/25:0) and 32.25 pmol PE P-Mix 3 (18:1p/15:0, 18:1p/19:0, 18:1p/25:0). Lipid standard preparations w done as described in^31^. Lipid extracts were resuspended in 60 µl methanol and samples were analyzed on a QTRAP 6500+ mass spectrometer (Sciex) with chip-based (HD-D ESI Chip, Advion Biosciences, USA) nano-electrospray infusion and ionization via a Triversa Nanomate (Advion Biosciences, Ithaca, USA) as previously described^31,32^. Data evaluation was done using LipidView (Sciex) and an in-house-developed software (ShinyLipids)

#### NOS enzymatic assay

Mammary glands were dissected and homogenized in NOS assay buffer and further processed following the Nitric Oxide Synthase Activity Assay kit (Abcam, ab211083) protocol for measuring enzymatic activity of nitric oxide synthase (NOS). Statistics were calculated by unpaired two-samples t-test following assessment of normality and equal variance using the Shapiro-Wilk’s test and F test, respectively.

#### Glycolysis inhibition experiments

Cells were seeded in 3D conditions (5µl of 80% Matrigel droplet) in black with clear flat bottom TC-treated imaging 96-well plates (Falcon, 353219). 3-bromopyruvate (Sigma-Aldrich, 16490) was added to the cell media, whereby following doses (µM) were tested: 250, 50, 25, 10, 2, 0 (vehicle, water). CellTox Green Cytotoxicity Assay (Promega, G8741) was performed according to manufacturer instructions to measure cell death after 72 hours or 48 hours of treatment. Green fluorescence was measured using EnVision plate reader (PerkinElmer) Resorufine/Amplex Red FP 535 for excitation and Europium 615 emission filter. Data analysis was performed using GraphPad Prism8, fold change was calculated by normalization to untreated control using transform function, transform Y values using Y = Y/K, different K for each dataset K = mean of untreated control for normal, tumor and residual separately. Experiments were reproduced two or three times; number of biological replicates is depicted in the figure legends and for each biological replicate there were 5-6 technical replicates take an average from technical replicates.

Images were taken over time-course of the experiment from using the high-throughput Olympus ScanR microscope in transmission mode. Each well of the 96-well plate was imaged using 1 ROI per with 21 Z-stacks (100 µm distance between stacks) at 4X magnification in a chamber with standard conditions (37 °C, 5% CO_2_). Projections of z-stacks and image stitching was done using a Fiji software.

#### DNA methylation profiles

After digestion of the Matrigel with collagenase and liberase, DNA was collected from the 3D cultures and extracted following Qiagen protocol for cultured cells (Qiagen Blood & Cell Culture DNA Mini kit, 13323). Bisulfite-free sequencing was performed at the Genomics Core Facility at EMBL Heidelberg.

Sequencing reads were aligned with the “qAlign” function of the R package “QuasR” version 1.22.1 ^33^ using the reference genome sequence for M. musculus from the R package “BSgenome.Mmusculus.UCSC.mm10” version 1.4.0 the R bowtie wrapper “Rbowtie” version 1.22.0 [Hahne F, Lerch A, Stadler MB. “Rbowtie: An R wrapper for bowtie and SpliceMap short read aligners.”] and^34^(rand the parameter setting modification: bisulfite=“undir”. Next, the DNA methylation was genome-wide quantified for all cytosine nucleotides in CpG context with the function “qMeth” from “QuasR” using the information from both strands combined. R V.3.5.1. (R Development Core Team) was used for conducting biostatistical analyses.

